# Genetic drift and genome reduction in the plant pathogen *Candidatus* Liberibacter solanacearum shapes a new enzyme in lysine biosynthesis

**DOI:** 10.1101/2023.04.17.537279

**Authors:** Jenna M. Gilkes, Rebekah A. Frampton, Amanda J. Board, André O. Hudson, Thomas G. Price, Deborah L. Crittenden, Andrew C. Muscroft-Taylor, Campbell R. Sheen, Grant R. Smith, Renwick C.J. Dobson

## Abstract

The effect of population bottlenecks and genome reduction on enzyme function is poorly understood. ‘*Candidatus* Liberibacter solanacearum’ is a bacterium with a reduced genome that is transmitted vertically to the egg of an infected psyllid—a population bottleneck that imposes genetic drift and is predicted to affect protein structure and function. Here, we define the effects of genome reduction and genetic drift on the function of *Ca*. L. solanacearum dihydrodipicolinate synthase (*C*LsoDHDPS), which catalyses the committed branchpoint reaction in diaminopimelate and lysine biosynthesis. We demonstrate that *C*LsoDHDPS is expressed in *Ca*. L. solanacearum and expression is increased ∼2-fold in the insect host compared to *in planta*. *C*LsoDHDPS has increased aggregation propensity, implying mutations have destabilised the enzyme but are compensated for through elevated chaperone expression and a stabilised oligomeric state. *C*LsoDHDPS uses a ternary-complex kinetic mechanism, which is unique among DHDPS enzymes, has unusually low catalytic ability, but an unusually high substrate affinity. Structural studies demonstrate that the active site is more open, and the structure of *C*LsoDHDPS with both pyruvate and the substrate analogue succinic-semialdehyde reveals that the product is both structurally and energetically different and therefore evolution has in this case fashioned a new enzyme. Our study reveals the effects of genome reduction and genetic drift on the function of essential enzymes and provides insights on bacteria-host co-evolutionary association. We suggest that bacteria with endosymbiotic lifestyles present a rich vein of interesting enzymes useful for understanding enzyme function and/or informing protein engineering efforts.

## Introduction

Bacteria evolved to occupy a broad range of niches that include extreme temperatures and pH fluxes. Some cultivate complex interactions with other organisms; for example, the obligate endosymbiotic associations bacteria have formed with plant phloem feeding insects (Nakabachi et al. 2006). Such interactions result in co-speciation of the bacteria endosymbiont and insect host that led to some bacteria reducing their genome size. This is thought to reflect the richness of host metabolites that allow for the loss of genes previously essential to the bacterial endosymbiont (Thao et al. 2000; Douglas 2006; Marais et al. 2008; Ibanez et al. 2014). The benefit of the relationship to the host insect is the acquisition of metabolic capabilities from the bacterial endosymbiont that are essential for the insect when feeding on nutrient limited sources, such as the phloem of plants (Wu et al. 2006).

*‘Candidatus* Liberibacter solanacearum’ is an unculturable, reduced genome (1.26 Mbp, *cf.* 5.6 Mbp for *Escherichia coli*), Gram-negative α-proteobacterium (Lin et al. 2011). It causes zebra chip disease in plants, a major problem in potato-growing areas causing significant economic impact on the potato industries in North and Central America, Europe and New Zealand (Secor et al. 2006; Munyaneza et al. 2007; Munyaneza 2010; Munyaneza 2012; Vereijssen et al. 2018). *Ca*. L. solanacearum persists in two diverse, but nutrient limiting niches: as an endosymbiont within an insect and as a pathogen within the phloem of an infected solanaceous plant. It is transmitted through these niches either horizontally, or vertically (**Figure 1**).

**Figure 1.**
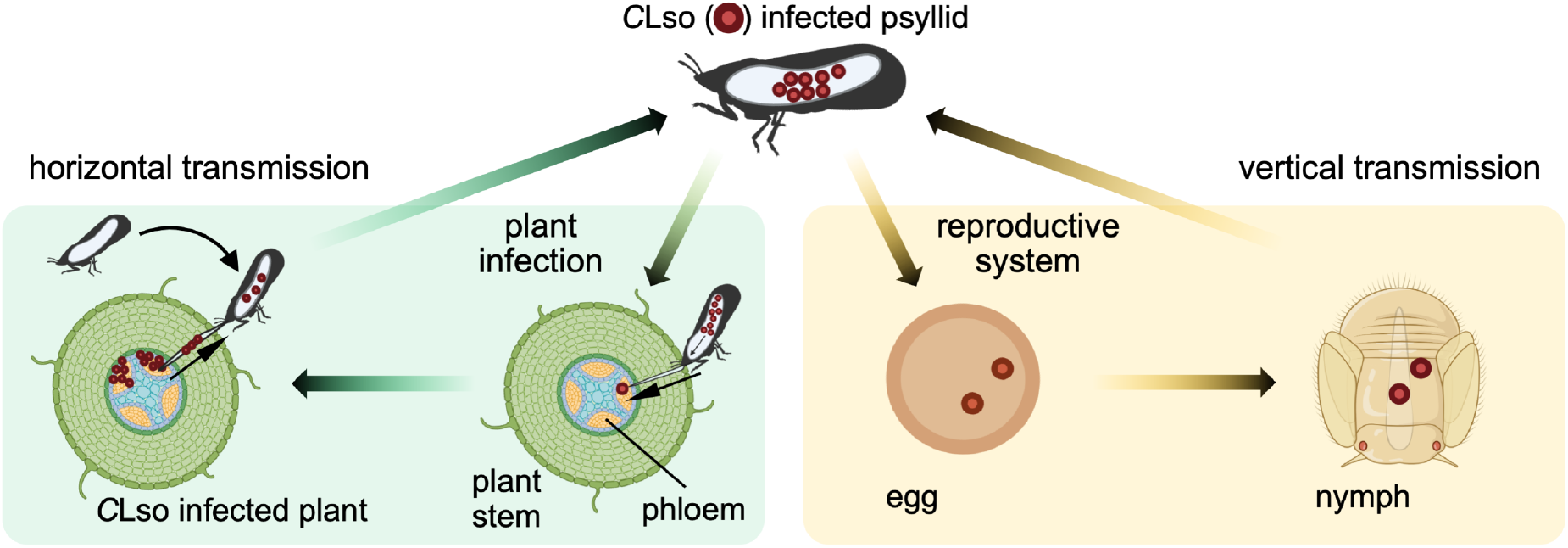
The lifecycle of *Ca. L. solanacearum* and its psyllid vector, *B. cockerelli.* Horizontal transmission, green box. A *Ca*. L. solanacearum infected psyllid feeds on the phloem sap of a health plant. *Ca*. L. solanacearum is transmitted *via* salivary secretions into a plant during psyllid feeding, which leads to *Ca*. L. solanacearum infection of the plant. A *Ca*. L. solanacearum free psyllid can then become infected upon feeding on the phloem of an infected plant. *Ca*. L. solanacearum within the gut can pass through the alimentary canal wall, move through the haemolymph to reach the salivary glands ready for reinfection. Vertical transmission, orange box. *Ca*. L. solanacearum are transmitted to the offspring (egg and nymph) of an infected female psyllid.

During horizontal transmission, the psyllid acquires the bacterium by feeding on infected solanaceous plants (**Figure 1**, green box). The bacterium is then vectored by the insect *Bactericera cockerelli* (Šulc) (Hemiptera: Triozidae), commonly known as the tomato potato psyllid (Munyaneza et al. 2007; Munyaneza 2010; Buchman et al. 2011; Buchman et al. 2012; Munyaneza 2012). To reinfect the plant, *Ca*. L. solanacearum crosses the insect alimentary canal wall and moves through the haemolymph to the salivary glands where the bacteria can then be transmitted to new host plants during psyllid feeding (Pitman et al. 2011; Cicero et al. 2017; Vereijssen et al. 2018).

During vertical transmission, bacteria are transmitted to the offspring of an infected female psyllid and multiply in the nymph and then adult insect (**Figure 1**, orange box) (Hansen et al. 2008; Casteel et al. 2012). Fluorescence *in-situ* hybridisation determined the spatial localisation of *Candidatus* Liberibacter asiaticus in the Asian citrus psyllid (Ammar et al. 2011), demonstrating near systemic infection the psyllid, including the ovaries. In addition, despite being reared on uninfected plant material, 3.6% of combined psyllid offspring (eggs, nymphs, and adults) derived from *Candidatus* Liberibacter asiaticus infected females tested positive for the bacterium (Pelz-Stelinski et al. 2010). Taken together, these reports support vertical transmission of *Candidatus* Liberibacter asiaticus in Asian citrus psyllid populations and it is likely that it also occurs with other Liberibacter species and their insect vectors (Pelz-Stelinski et al. 2010; Ammar et al. 2011; Kelley and Pelz-Stelinski 2019).

Vertical transmission results in a reduction in the population of bacteria transmitted to the insect offspring, a population bottleneck in sustaining symbionts through successive generations. For bacterial endosymbionts with lifestyles such as *C*Lso, there are consequences for their genome and protein complement when compared to free living bacteria. Successive population bottlenecks allow deleterious mutations to become fixed through genetic drift, since natural selection is limited (Wernegreen 2002; McCutcheon and Moran 2012; Wernegreen 2015). Secondly, a consequence of genome reduction is the degradation of genes for DNA repair and a bias towards adenine and thymine nucleotides (Kelkar and Ochman 2013). This translates to a mutational bias towards hydrophobic residues at the protein level that results in the destabilisation of protein structure (Kelkar and Ochman 2013). Furthermore, the fixation of deleterious mutations by genetic drift affects the thermodynamic stability of proteins (Ham et al. 2003), destabilises protein secondary structure and makes the proteins prone to misfolding and aggregation (Ham et al. 2003). For *C*Lso, horizontal transmission (*via* the plant) likely offsets some of the consequences of vertical transmission. Here, we investigate the effect of population bottlenecks and genome reduction on enzyme function.

*Ca*. L. solanacearum has lost many of the enzymes required for *de novo* amino acid biosynthesis, but is predicted to retain the genes for functional enzymes that synthesise lysine (Lin et al. 2011), suggesting this pathway is essential. *Ca*. L. solanacearum synthesises lysine *via* the diaminopimelate pathway, named after the immediate precursor of lysine, *meso*-diaminopimelate, which is a major constituent of the bacterial peptidoglycan cell wall in Gram-negative bacteria (Cox et al. 2000; Hutton et al. 2007) (**Figure 2A**). Thus, compounds that inhibit the diaminopimelate pathway could be a good strategy to control bacterial infections (Girodeau et al. 1986; Cox 1996; Cox et al. 2000; Turner et al. 2005; Hutton et al. 2007; Boughton et al. 2008; Mitsakos et al. 2008; Christoff et al. 2019; Costa et al. 2021), including *Ca*. L. solanacearum infections in plants.

**Figure 2.**
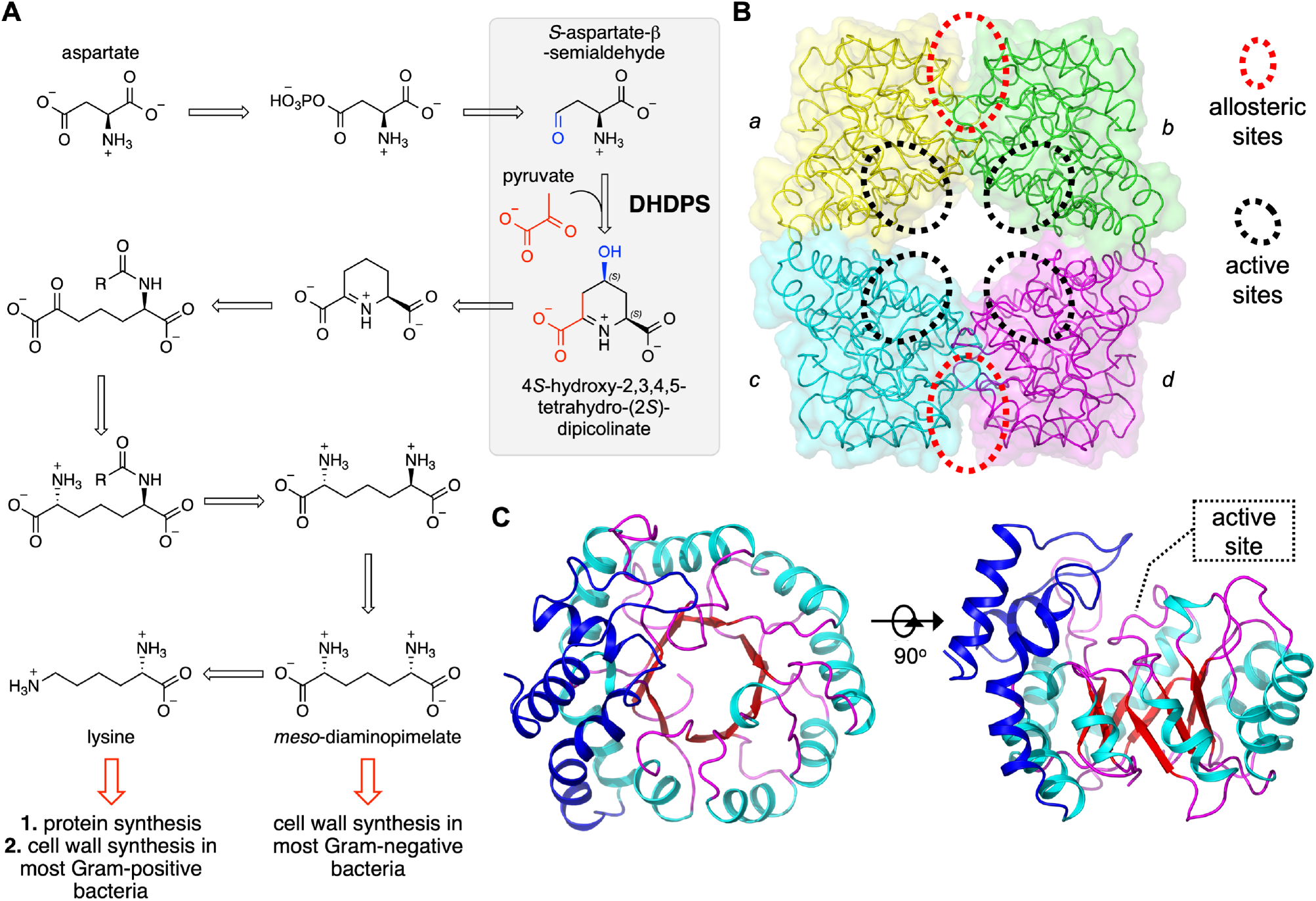
The diaminopimelate pathway and structure of DHDPS. **A.** The diaminopimelate pathway that affords precursors for cell wall and protein synthesis. Highlighted in the grey box is the reaction catalysed by DHDPS—the condensation of pyruvate and *S*-aspartate-β-semialdehyde catalysed to produce (4*S*)-hydroxy-2,3,4,5- tetrahydro-(*2S*)-dipicolinate. **B.** The tetrameric structure of the *Ec*DHDPS (PDB ID: 1YXC) with each of the monomers (*a*, *b*, *c* and *d*) differentially coloured. The tight-dimer interface is formed between monomers *a* and *b*, as well as between monomers *c* and *d*. The weak-dimer interface is formed between monomers *a* and *c*, as well as between *b* and *d*. **C.** The TIM barrel structure of the DHDPS monomer, highlighting the location of the active site within the TIM barrel.

Dihydrodipicolinate synthase (abbreviated to DHDPS) catalyses the condensation of pyruvate and *S*-aspartate-β-semialdehyde to form (4*S*)-hydroxy-2,3,4,5-tetrahydro-(2*S*)- dipicolinate, and this is the first committed step of lysine biosynthesis in the diaminopimelate pathway (Blickling, Renner, et al. 1997; Dobson, Valegård, et al. 2004; Burgess et al. 2008; Kefala et al. 2008; Domigan et al. 2009) (**Figure 2A**). We note the explicit stereochemistry of the (4*S*)-hydroxy position, which becomes important later. In plants and Gram-negative bacteria, DHDPS is also allosterically feedback inhibited by lysine, the end-product of the pathway, and therefore a key point of regulation (Dobson, Griffin, et al. 2004; Dobson et al. 2005; Atkinson et al. 2012). The structure of DHDPS has been extensively studied and usually comprises a homotetramer of (β/α)_8_-barrel monomers that arrange into a dimer of ‘tight-dimers’ (**Figure 2B**). The active site is located in the centre of a (β/α)_8_-barrel in each monomer (**Figure 2C**), while the allosteric lysine-binding site is situated distal from the active site in the cleft at the tight-dimer interface, with one lysine binding per monomer, but each lysine molecule being co-ordinated by residues from each monomer within the tight-dimer.

The recent development of a method to isolate and characterise recombinant proteins derived from *Ca*. L. solanacearum (Gilkes et al. 2019) provides a unique opportunity to evaluate the evolutionary consequences of highly specialised host-dependent lifestyles, including understanding the effects of genetic drift and genome evolution on the structure and function of enzymes from reduced genome organisms. Furthermore, because DHDPS has been extensively studied from many bacteria, this enzyme provides an ideal model system for defining the effects of genome reduction and genetic drift on protein structure and enzyme function. Here, we demonstrate that the lifestyle of *Ca*. L. solanacearum has resulted in a DHDPS homologue with highly unusual functional properties, but with surprisingly faithful structural conservation.

## Results and Discussion

### CLsoDHDPS is expressed preferentially within the psyllid, compared to the plant

We first verified that the lysine biosynthetic enzyme dihydrodipicolinate synthase (DHDPS) is expressed in *C*Lso. Bacteria adapt to their immediate environment by optimising their gene expression patterns in response to signals from that environment (Yan et al. 2013): expression of genes necessary in a given medium is up-regulated (activated), while those that are unnecessary are down-regulated (repressed) (Chowdhury et al. 1996). As such, defining gene expression helps to identify those genes necessary for *Ca*. L. solanacearum survival within either the psyllid or plant host. We tested whether the *Ca*. L. solanacearum *dapA* gene, which encodes the enzyme DHDPS, is expressed by the bacterium in native settings (tomato plant and psyllid host) using quantitative reverse transcription-polymerase chain reaction (qRT-PCR) to test for the presence of *Ca*. L. solanacearum *dapA* mRNA (**Figure 3A**). In both environments, we find high expression levels of *Ca*. L. solanacearum *dapA* mRNA relative to the housekeeping genes *Ca*. L. solanacearum *recA* (recombinase A) and *Ca*. L. solanacearum *rpb* (DNA-directed RNA polymerase beta subunit), evidence that *dapA* is indeed transcribed. Moreover, the expression of *Ca*. L. solanacearum *dapA* is increased ∼2-fold in the insect host compared to *in planta,* again relative to each housekeeping gene, *recA* and *rpb.* Thus, *dapA* is expressed in *Ca*. L. solanacearum and the level of expression differs when *Ca*. L. solanacearum is in the psyllid gut compared to the phloem of the plant. The physiological reason for the differential expression in the two environments is unknown, but we speculate that upregulation of lysine biosynthesis when in the gut may reflect increased growth rate and therefore higher needs of *meso-* diaminopimelate for the bacterial cell wall and lysine for protein production.

**Figure 3.**
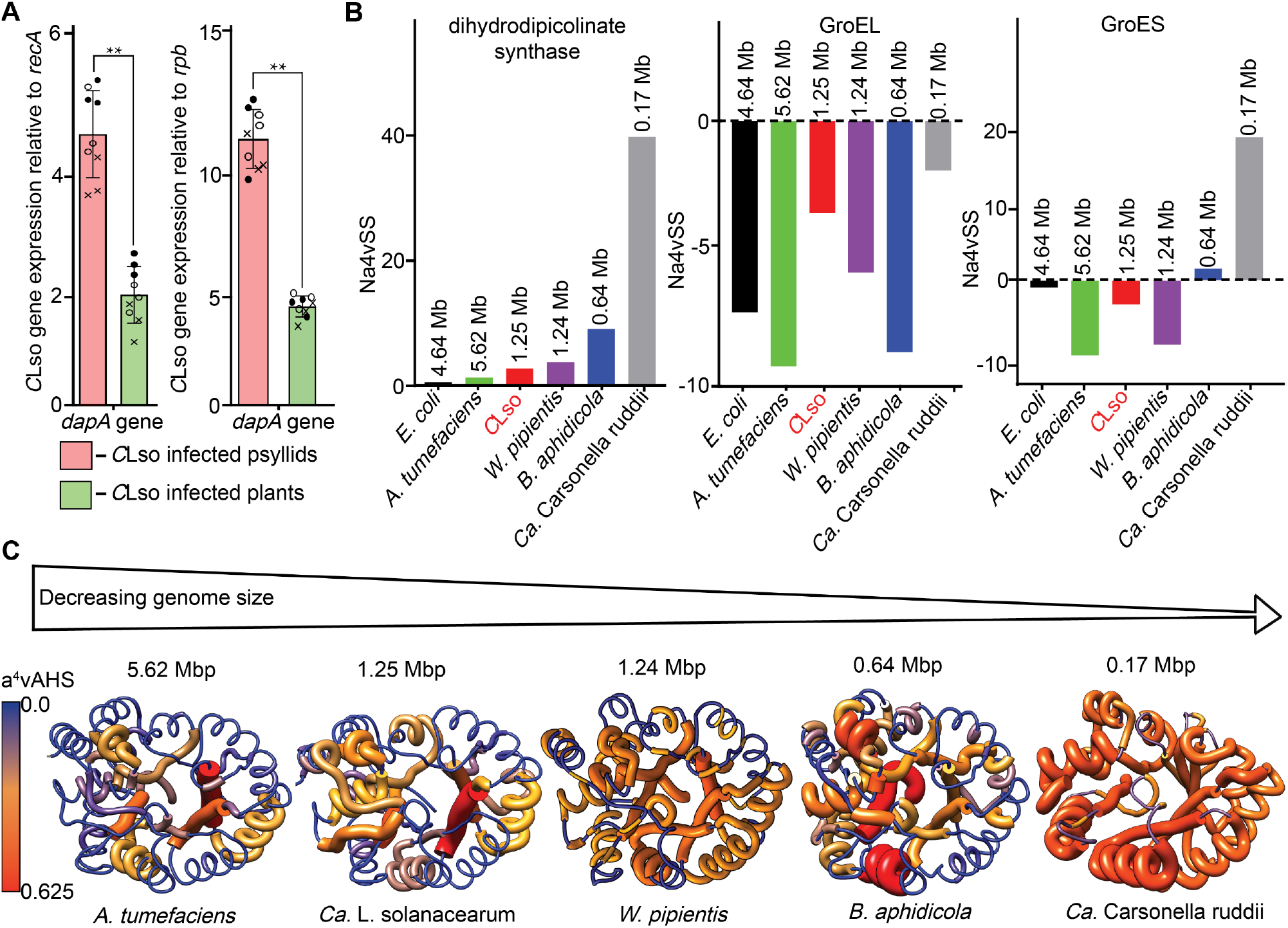
*DapA* Gene expression *in psyllids* and *in planta* and aggregation analysis of *Ca.* L. solanacearum proteins. **A.** Level of gene expression determined using qRT-PCR in relation to the housekeeping genes *recA* (recombinase A, KJZ80672, DJ66_1282) and *rpb* (DNA-directed RNA polymerase beta subunit, KJZ81364.1, DJ66_1258). Mean ± standard deviation for three biological (o, ●, **×**) and three technical replicates are plotted. Asterisks indicate statistically significant differences in expression levels between *Ca*. L. solanacearum infected psyllids and plants (*P ≤ 0.05, **P ≤ 0.01; *t*-test; n=3). Fold change (log_2_) is the relative gene expression in psyllid versus *in planta* for *C*Lso. Positive values indicate the gene was overexpressed in psyllids compared with *in planta*. The relative gene expression of *dap*A is increased 1.45 ± 1.15 fold (P-value = 0.004) in the psyllid compared to in the plant when normalised to *recA* and 3.2 ± 0.52 fold (P-value = 0.008) when normalised to *rpb*. **B** shows the aggregation propensity analysis of DHDPS and chaperones GroEL and GroES determined by AGGRESCAN analysis for various organisms. **C** shows the increase in protein aggregation propensity of the DHDPS monomer with decreasing genome size. The a^4^vAHS is the maximum aggregation propensity value. The structures of the *Wolbachia pipientis*, *B. aphidicola* and *Ca*. Carsonella ruddii DHDPS in the image were generated using SwissModel, whilst the *A. tumefaciens* structure was sourced from the PDB (4I7U) and *C*LsoDHDPS from the structure solved in this study.

### CLsoDHDPS is predicted to have an increased aggregation propensity

The transition of free-living *Ca*. L. solanacearum to an endosymbiont with an insect led to genome reduction (Lin et al. 2011) and population bottlenecks in the lifecycle (Hansen et al. 2008; Ammar et al. 2011; Casteel et al. 2012; Workneh et al. 2016). These are predicted to have a general destabilising effect on protein structure and increase the propensity for aggregation, which in turn affects function. We tested whether *C*LsoDHDPS is destabilised compared with homologues from other reduced genome bacteria and free-living, mesophilic bacteria.

To determine whether the *Ca*. L. solanacearum proteins have a greater propensity for aggregation, AGGREGSCAN (Conchillo-Solé et al. 2007) was used to predict aggregation-prone regions in the protein sequences of interest. AGGRESCAN is based on an aggregation-propensity scale for natural amino acids and assumes that short and specific sequence stretches modulate protein aggregation (Conchillo-Solé et al. 2007). The intrinsic aggregation properties for DHDPS homologues and the chaperone proteins GroEL and GroES (discussed in the next section) were predicted and compared across *Escherichia coli* (PDB id 1YXC) and *Agrobacterium tumefaciens* (PDB id 4I7U), as well as the reduced genome organisms of *Ca*. L. solanacearum (KJZ81861), *Wolbachia pipientis* (KLT22486.1), *Buchnera aphidicola* (BAB12815.1) and *Candidatus* Carsonella ruddii (AGS06617.1).

The average aggregation propensity of the sequence (Na^4^vSS) for *C*LsoDHDPS is shown in **Figure 3B**. DHDPS from the reduced genome organisms *B. aphidicola*, *Ca.* Carsonella ruddii and *Ca*. L. solanacearum have a greater propensity for protein aggregation when compared against *E. coli* and *A. tumefaciens* (**Figure 3B**)—here, an increased positive Na^4^vSS score means an increase in the aggregation propensity for the protein. When we broaden the analysis to enzymes within the lysine biosynthetic pathway, we see the same trend (**Supplementary Figure i**). The aggregation propensity for *C*LsoDHDPS is also demonstrated in **Figure 3C**, which maps average hot spot area (a^4^vAHS) onto the tertiary structure of the DHDPS enzymes and shows there is an increase in a^4^vAHS as genome size decreases. The *B. aphidicola* and *Ca.* Carsonella ruddii DHDPS enzymes are shown to be most prone to aggregation, in line with the computational studies (Ham et al. 2003), which suggest that *B. aphidicola* proteins have a reduced and slower protein folding efficiency, increased misfolding and aggregation propensity, or are unstable in their native conformation. Overall, the results are consistent with an increase in protein instability as genome size decreases.

We next examined and compared the *C*LsoDHDPS protein sequence with that of other bacterial and plant homologues (**Supplementary Figure ii**). In general, there are no obvious sequence differences when comparing *C*LsoDHDPS or homologues from other genome reduced species with well-established bacterial DHDPS systems. For example, the *C*LsoDHDPS is similar in length (*i.e.*, no deletions or insertions) and generally similar across the sequence (*i.e.*, no areas of obvious difference that map to regions with increased aggregation).

### CLsoDHDPS chaperone proteins GroEL and GroES showed a low propensity for aggregation

The effects of genetic drift and genome reduction on bacterial endosymbionts should lead to an eventual decline in biological fitness and eventual extinction in the absence of recombination, however, the stable lifestyle of endosymbionts suggests a compensating mechanism (Fares, Barrio, et al. 2002; Fares, Ruiz-González, et al. 2002; Fares et al. 2004). The retention and increased abundance of chaperone proteins in endosymbionts is thought to assist protein folding and rescue non-functional protein conformations (Fares, Ruiz-González, et al. 2002; Fares et al. 2004; Tokuriki and Dan S. Tawfik 2009). Importantly, chaperones are abundantly expressed and evolutionary conserved among intracellular bacteria (Fares, Ruiz-González, et al. 2002; Fares et al. 2004; Tokuriki and Dan S. Tawfik 2009; Nachappa et al. 2012).

We previously demonstrated that overexpression and purification of folded and functional recombinant *Ca*. L. solanacearum enzymes, including *C*LsoDHDPS, required the concurrent overexpression of *E. coli* chaperone proteins GroEL and GroES (Gilkes et al. 2019). Moreover, others have shown that the expression of functional *E. coli* DHDPS (*Ec*DHDPS) is dependent on the GroEL/GroES chaperone system—disrupting the GroEL/GroES system results in *E. coli* lysis because DHDPS cannot fold and therefore the cell wall constituent diaminopimelate cannot be synthesized (McLennan and Masters 1998). In the *E. coli* setting, GroEL/ES accelerates the folding of DHDPS 30-fold by catalysing segmental structure formation in the TIM-barrel and lowering the entropy of the energy barrier (Georgescauld et al. 2014). Thus, the requirement to co-express *E. coli* chaperone proteins to high levels with the *C*LsoDHDPS could indicate the instability of the protein when compared to *Ec*DHDPS, which only requires chromosomal chaperonin expression levels when overexpressed.

It is thought that the abundant overexpression of chaperones in *Ca*. L. solanacearum (Nachappa et al. 2012) assist the folding of conformationally damaged proteins that occurs as a result of genome reduction and genetic drift (Fares et al. 2004; Fares et al. 2005). Included in **Figure 3B** is the aggregation propensity for *Ca*. L. solanacearum GroEL and GroES, which suggests that these proteins have decreased aggregation propensity in contrast to the *Ca*. L. solanacearum enzymes of lysine biosynthesis. That is, they retain stability and function. We hypothesise that these *Ca*. L. solanacearum chaperonins are buffering against mutational effects and are subject to purifying selection rather than genetic drift, as they are required for stabilising and folding of *Ca*. L. solanacearum proteins (Moran 1996; Fares, Barrio, et al. 2002; Fares, Ruiz-González, et al. 2002; Rutherford 2003). The GroEL chaperone from all organisms tested showed a low propensity for aggregation. This is consistent with the proposal that *Ca*. L. solanacearum chaperone proteins buffer against the deleterious mutations that accumulate under strong genetic drift from the bacterium’s intracellular lifecycle and subsequent fitness decline (Moran 1996; Fares, Barrio, et al. 2002; Fares, Ruiz-González, et al. 2002; Rutherford 2003). Interestingly, biological systems with high GroEL/GroES overexpression increased the number of and buffered against the accumulation of mutations that are destabilising and increased enzymatic variability (Tokuriki and Dan S. Tawfik 2009; Tokuriki and Dan S Tawfik 2009).

Overall, we demonstrate that *C*LsoDHDPS is likely to be more prone to aggregation compared to *E. coli* and *A. tumefaciens,* and more similar in aggregation propensity to other reduced genome bacteria. We also suggest that the GroEL/ES chaperone systems are stable and functional, consistent with the proposal that this is the mechanism by which genome reduced bacteria overcome deleterious mutations. As such, not only is DHDPS expressed in *Ca*. L. solanacearum, but is likely to be in a folded and functional form, facilitated by the GroEL/ES chaperone system.

### CLsoDHDPS uses the ternary-complex kinetic mechanism, but retains allosteric inhibition by lysine

We tested whether genome reduction in *Ca*. L. solanacearum had driven the evolution of altered catalytic functions in *C*LsoDHDPS. Such differences in function could be exploited for designing effective antibacterial agents that target DHDPS, especially if they block an active site that is significantly different to endogenous plant or other bacterial DHDPS enzymes. Recombinant *C*LsoDHDPS was expressed and purified using the method previously described (Gilkes et al. 2019). Initial rate kinetic data were collected using an established assay (Dobson, Valegård, et al. 2004), varying the concentration of both substrates, pyruvate and *S*-aspartate-β-semialdehyde. As a first step, we determined the kinetic mechanism by which *C*LsoDHDPS operates.

Surprisingly, *C*LsoDHDPS follows the ternary-complex kinetic mechanism (**Figure 4A**, top), whereas all DHDPS enzymes characterised to date follow the ping-pong kinetic mechanism (bottom), which suggests that the order of substrate binding and product release has changed and therefore catalytic mechanism is different in this homologue. Initial rate data for varied pyruvate and *S*-aspartate-β-semialdehyde concentrations fit the ternary-complex model with an R^2^ value of 0.99 (**Figure 4B** and **Table 1**). The Akaike test was used to determine the best fit between the ping-pong and ternary-complex models, and estimated that the probability of the ping-pong model being correct was <0.01%, whereas the probability of the ternary-complex model was 99.99% with a difference in the AIC value of 18.9. Unlike the ping-pong kinetic model, the ternary-complex kinetic model can be either ordered or random with respect to substrate binding and product release. However, assuming that the Schiff base/enamine needs to form first to enable C-C bond formation in the *C*LsoDHDPS catalytic mechanism, we propose that pyruvate is the first substrate to bind and that H_2_O and H^+^ are the first products released.

**Figure 4.**
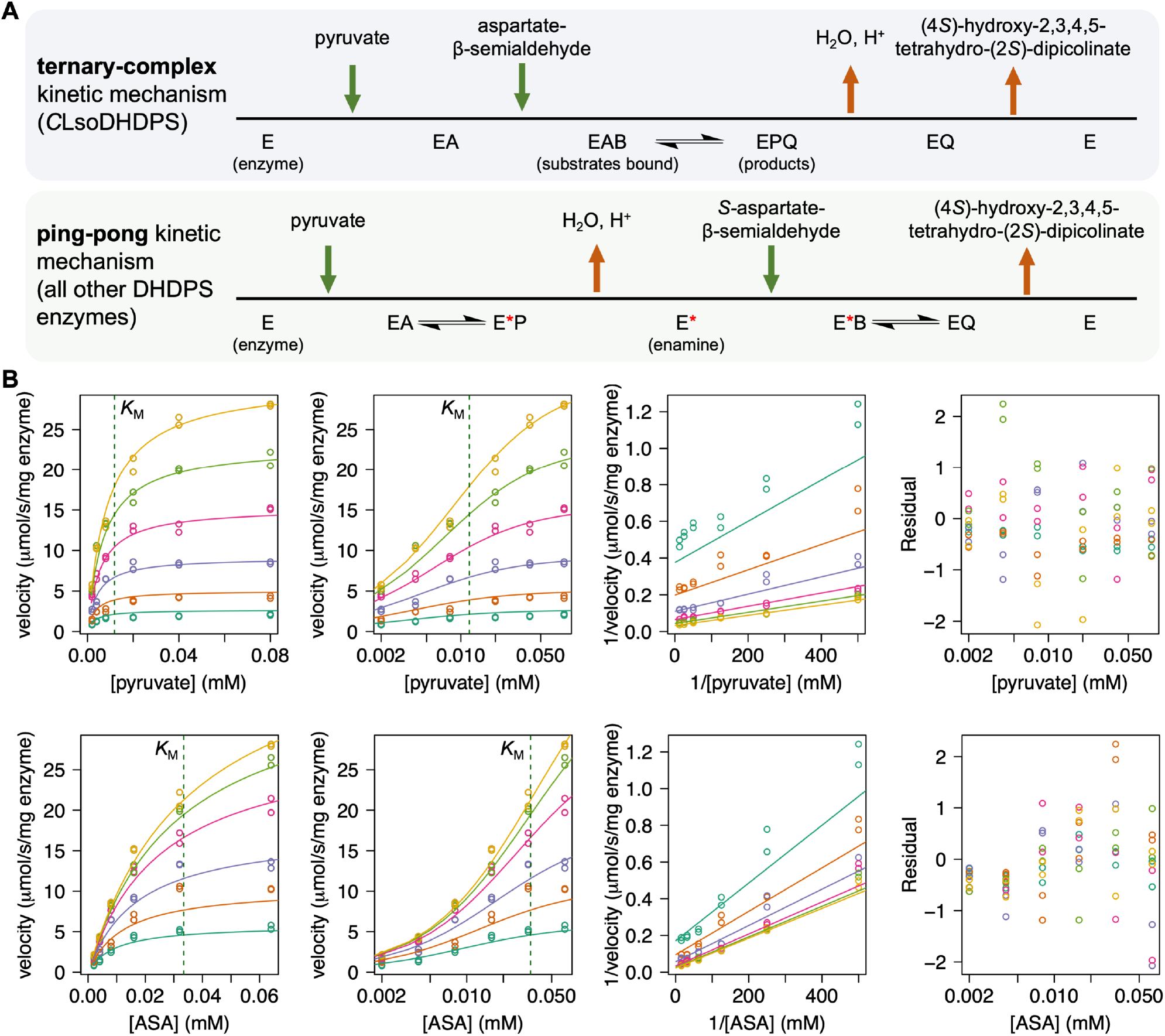
Enzyme kinetic analysis of *C*LsoDHDPS. **A.** Schemes for the ternary-complex and the ping-pong kinetic models. They differ in the order of substrate binding and product release. **B.** Initial rate data best fit to the ternary-complex kinetic model. The data has an R^2^ value of fit is 0.99. The plots show rates with respect to pyruvate (top) and *S*-aspartate-β- semialdehyde (bottom), and were generated using R following non-linear regression of the data against the ternary-complex model. The four plots from left to right are the direct plot, the direct plot with the substrate concentration on the log scale, the Lineweaver Burk plot, and the residuals of the fit. For the plots with respect to pyruvate (top), the *S*-aspartate-β- semialdehyde concentrations are O = 0.064 mM, O = 0.032 mM, O = 0.016 mM, O = 0.008 mM, O = 0.004 mM, and O = 0.002 mM. For the plots with respect to *S*-aspartate-β- semialdehyde (*S*-ASA) (bottom), the pyruvate concentrations are O = 0.08 mM, O = 0.04 mM, O = 0.02 mM, O = 0.008 mM, O = 0.004 mM, and 0 = 0.002 mM. The green dotted line shows the fitted *K*_M_ values for pyruvate (top) and *S*-aspartate-β-semialdehyde (bottom).

**Table 1.**
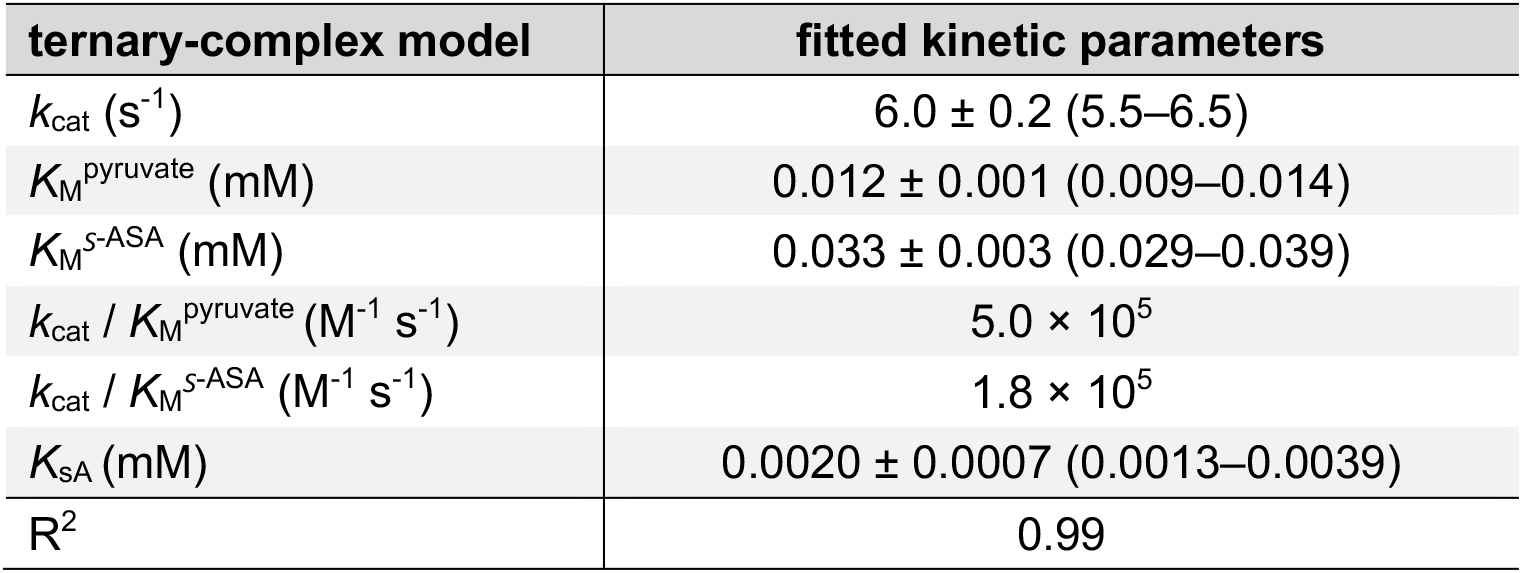
Kinetic parameters for *C*LsoDHDPS with respect to pyruvate and *S*-aspartate-β-semialdehyde (*S*-ASA). The data were fit to ternary-complex model using R, as in **Figure 4**. The standard error from the fit for each parameter is shown and the 95% confidence intervals are in brackets.

| ternary-complex model | fitted kinetic parameters |
| --- | --- |
| $k_{\text{cat}}$ ($\text{s}^{-1}$) | $6.0 \pm 0.2$ (5.5–6.5) |
| $K_{\text{M}}^{\text{pyruvate}}$ (mM) | $0.012 \pm 0.001$ (0.009–0.014) |
| $K_{\text{M}}^{\text{S-ASA}}$ (mM) | $0.033 \pm 0.003$ (0.029–0.039) |
| $k_{\text{cat}} / K_{\text{M}}^{\text{pyruvate}}$ ($\text{M}^{-1} \text{s}^{-1}$) | $5.0 \times 10^5$ |
| $k_{\text{cat}} / K_{\text{M}}^{\text{S-ASA}}$ ($\text{M}^{-1} \text{s}^{-1}$) | $1.8 \times 10^5$ |
| $K_{\text{SA}}$ (mM) | $0.0020 \pm 0.0007$ (0.0013–0.0039) |
| $R^2$ | 0.99 |

The kinetic parameters for *C*LsoDHDPS are also surprising. Firstly, the Michaelis constants (*K*_M_) for the substrates were 0.012 ± 0.001 mM with respect to pyruvate and 0.033 ± 0.003 mM with respect to *S*-aspartate-β-semialdehyde—these are considerably lower when compared to the Michaelis constants of other bacterial homologues (**Supplementary Table i**) and suggest the enzyme has a high affinity for its substrates. Secondly, the turnover number (*k*_cat_) is 6.0 ± 0.2 s^-1^ and is also considerably lower when compared to other bacterial homologues—the only comparable homologue is from *Thermotoga maritima*, but in that case the rates were determine far below the optimal temperature for this enzyme. Since *C*LsoDHDPS uses a ternary-complex kinetic mechanism, we also report a true substrate dissociation constant for pyruvate (*K*_sA_) of 0.0020 ± 0.0007 mM. Despite the low turnover number, the specificity constants are comparable with other bacterial homologues (**Supplementary Table i**), driven by improved affinity for the substrates (lower *K*_M_ values). One interpretation is that the accumulation of mutations in *C*LsoDHDPS during genome reduction may have led to a trade-off between the declining ability to catalyse the chemical transformation, but improved affinity for the substrates.

Next, we tested whether *C*LsoDHDPS retains feedback inhibition by the pathway end product, lysine, as seen in other DHDPS homologues studied to date (Dobson, Griffin, et al. 2004; Rice et al. 2008; Domigan et al. 2009). To test this, initial rate data were collected as above, but with varying concentrations of lysine. The initial rate for the enzyme decreases when titrating increasing concentrations of lysine into the assay, demonstrating that *C*LsoDHDPS retains its ability to be allosterically inhibited by lysine (**Supplementary Figure iii**). In addition, lysine acts as a partial inhibitor, since at saturating lysine concentrations some activity is retained, a behaviour that is also seen in *Ec*DHDPS (Dobson, Griffin, et al. 2004). Inhibition of *C*LsoDHDPS by lysine when pyruvate is the varied substrate displayed mixed model inhibition with a *K*_ic_ of 0.37 ± 0.07 mM and *K*_iu_ of 0.121 ± 0.005 mM (**Table 2**). Inhibition of *C*LsoDHDPS by lysine when *S*-aspartate-β-semialdehyde is the varied substrate displayed partial uncompetitive inhibition with a *K*_iu_ of 0.27 ± 0.02 mM (**Table 2**).

**Table 2.** Kinetic parameters for lysine inhibition of CLsoDHDPS with respect to pyruvate and S-aspartate-β-semialdehyde as a substrate. The data were fit to respective kinetic models using R, as in **Supplementary Figure iii**. The standard error of the fit for each parameter is shown and the 95% confidence intervals are in brackets.

|  | fitted kinetic parameters |  |
| --- | --- | --- |
| varied substrate | pyruvate | S-aspartate-β-semialdehyde |
| kinetic model | mixed inhibition | partial uncompetitive inhibition |
| $K_{\text{M}}$ (mM) | $0.0110 \pm 0.0005$ (0.010–0.012) | $0.022 \pm 0.001$ (0.020–0.024) |
| $K_{\text{ic}}$ (mM) | $0.37 \pm 0.07$ (0.26–0.59) | - |
| $K_{\text{iu}}$ (mM) | $0.121 \pm 0.005$ (0.112–0.131) | $0.27 \pm 0.02$ (0.23–0.31) |
| $V_{\text{max}}$ ($\mu\text{mol}/\text{min}/\text{mg}$) | $32.4 \pm 0.5$ (31.4–33.4) | $33.1 \pm 0.8$ (31.5–34.7) |
| $V_{\text{max}2}$ ($\mu\text{mol}/\text{min}/\text{mg}$) | - | $1.0 \pm 0.3$ (0.5–1.5) |
| $R^2$ for the fit | 0.99 | 0.98 |

Considering our kinetic experiments together and despite predicting an increased propensity for aggregation, we verify that *C*LsoDHDPS is active, suggesting that although *Ca*. L. solanacearum has lost many of its biosynthetic genes, it has retained lysine biosynthesis. We demonstrate that *C*LsoDHDPS uses a different kinetic mechanism to other DHDPS enzymes, which implies that the catalytic mechanism may be altered in some way, including the order of substrate binding and product release. We also find that the *K*_M_ values for the substrates are considerably lower compared to other homologues, reflecting a higher affinity for the substrates. Although kinetically unusual, *C*LsoDHDPS retains allosteric inhibition by lysine.

### Crystal structures of CLsoDHDPS demonstrate a conserved structure, but larger tetrameric interfaces

Next, we solved a series of ligand bound *C*LsoDHDPS structures using crystallography to test whether evolutionary pressures resulted in structural changes that account for the altered kinetic behaviour. We determined and compared the following X-ray crystal structures: 1) with the allosteric inhibitor lysine, PDB id 7LVL; 2) with the substrate pyruvate, PDB id 7LOY; and 3) with pyruvate and succinic semi-aldehyde, an analogue of the substrate *S*-aspartate-β-semialdehyde, PDB id 8GEK. We had difficulty crystallising the structure of apo-*C*LsoDHDPS, perhaps reflecting some unknown structural heterogeneity that prevented crystal formation. Nonetheless, each of these structures were determined to atomic resolution (1.93–2.40 Å), with sensible geometry and *R*_free_ values (0.209–0.235) (**Supplementary Table iii**). Below, we outline the broad similarities and differences with homologous DHDPS enzymes, before focusing on the active and allosteric sites of the ligand bound structures in later sections.

We first examined the oligomeric structure of *C*LsoDHDPS, since DHDPS enzymes can form tetrameric (Dobson et al. 2005; Domigan et al. 2009; Atkinson et al. 2018) and/or dimeric structures (Burgess et al. 2008; Atkinson et al. 2018). The oligomeric state is known to be important for function because both the active site and the allosteric binding site comprise residues from the two monomers that form the tight-interface (the yellow and green monomers in **Figure 2B**) (Griffin et al. 2008; Pearce et al. 2011). The asymmetric unit for each *C*LsoDHDPS crystal structure contained six monomers that form the typical bacterial tetrameric structure (symmetry operations are needed to complete the second tetramer) (**Figure 5A**). The tetrameric structure is in agreement with sedimentation velocity experiments that demonstrate that unliganded *C*LsoDHDPS is tetrameric in solution (Gilkes et al. 2019). A structural alignment of *C*LsoDHDPS with other tetrameric bacterial DHDPS homologues from *E. coli*, *Thermotoga maritima*, *Campylobacter jejuni, A. tumefaciens* and *Bacillus anthracis* demonstrates that the *A. tumefaciens* DHDPS tetramer has the closest overall structural similarity, with the lowest root mean square deviation of 1.0 Å (α-carbons, **Figure 5B**).

**Figure 5.**
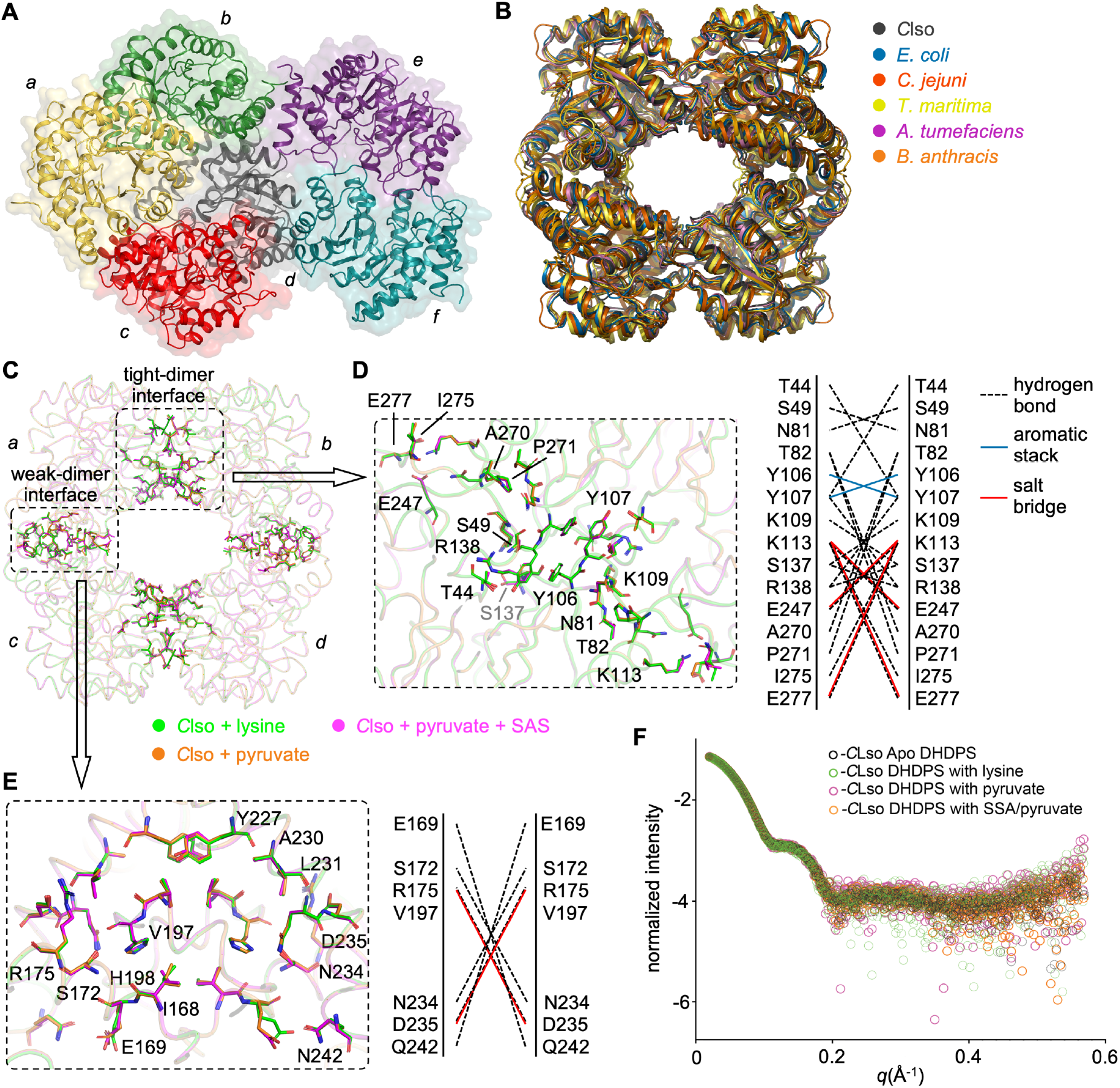
Structure of the *C*LsoDHDPS. **A.** The position of the six monomers in the *C*LsoDHDPS asymmetric unit. **B.** The *C*LsoDHDPS tetramer (black, PDB ID: 7LOY) aligned with the DHDPS tetramers for *E. coli* (blue, PDB ID: 1YXC, r.m.s.d. = 1.2), *Campylobacter jejuni* (red, PDB ID: 4R53, r.m.s.d. = 1.8), *T. maritima* (yellow, PDB ID: 3PB2, r.m.s.d. = 2.2), *A. tumefaciens* (magenta, PDB ID: 4I7U, r.m.s.d. = 1.0) and *B. anthracis* (orange, PDB ID: 3HIJ, r.m.s.d. = 1.1). Monomers are indicated by *a*/*b*/*c*/*d*, as in Figure 2B. **C.** An overlay of the liganded *C*LsoDHDPS tetramers. The *C*LsoDHDPS + lysine structure (green) is overlaid with the *C*LsoDHDPS + pyruvate (yellow, r.m.s.d. = 0.14) and the *C*LsoDHDPS + SSA + pyruvate (magenta, r.m.s.d. = 0.16) structures. The structures suggest that ligand binding does not alter the structure of the active and allosteric sites as there is no overall conformational change in the structure (which is confirmed in solution using small angle X-ray scattering in **F**). The position of the residues at the tight-dimer interface between monomers *a* and *b (c* and *d)* and the residues at the weak dimer interface between monomers *a* and *c* (*b* and *d*) are shown. **D.** and **E.** focus of the tight dimer and weak dimer interfaces, respectively, as determined by PISA. Broken lines indicate hydrogen-bonding, red lines indicate salt bridges, and blue lines aromatic stacking. **F.** Small angle scattering data comparing the apo *C*LsoDHDPS with ligand bound *C*LsoDHDPS demonstrates that ligand binding does not alter the structure of the enzyme in solution (**O** = apo *C*LsoDHDPS, **O** = *C*LsoDHDPS + lysine, O = *C*LsoDHDPS + pyruvate, and O *C*LsoDHDPS + SSA + pyruvate).

To determine whether binding either the substrates (pyruvate and succinic semi-aldehyde) or the allosteric inhibitor (lysine) result in subtle conformational changes across the *C*LsoDHDPS tetramer, the ligand bound tetrameric structures were overlayed and changes at the interfaces were investigated (**Figure 5C**). We note that in all the *C*LsoDHDPS structures presented, the nature of both the tight-dimer and weak-dimer interfaces do not change, suggesting that ligand binding does not change the orientation of the monomers within the tetramer, as compared for example with O_2_ binding to haemoglobin.

The tight-dimer interface between the *a/b* (and *c/d*) monomers includes residues Tyr106 and Tyr107 that are required for the interlocking of the monomers and contribute to the active site in the opposing monomer (**Figure 5D**). The Protein Interfaces, Surfaces and Assemblies (PISA) program (Krissinel and Henrick 2007) as used to calculate the buried surface area at the tight-dimer interface of *C*LsoDHDPS, which is on average ∼1520 Å^2^ (BSA in **Table 3**) with ∼20 hydrogen bonds and five salt bridges (**Figure 5D**). Compared to other homologues, *C*LsoDHDPS has considerably more contacts (hydrogen bonds and salt bridges) at this interface and it buries a ∼10% larger surface area from solvent.

**Table 3.** Comparison of the tight-dimer and weak-dimer interfaces of *C*LsoDHDPS (as shown in **Figure 5C**) and other characterised DHDPS enzymes using PISA (Krissinel and Henrick 2007). Succinic semi-aldehyde is abbreviated to SSA and buried surface area is abbreviated to BSA.

| Organism | Ligand | PDB ID | Tight-dimer interface<br>(monomers *a/b*, *c/d*) |  |  | Weak-dimer interface<br>(monomers *a/c*, *b/d*) |  |  |
| --- | --- | --- | --- | --- | --- | --- | --- | --- |
|  |  |  | BSA<br>(Å<sup>2</sup>) | hydrogen<br>bonds | salt<br>bridges | BSA<br>(Å<sup>2</sup>) | hydrogen<br>bonds | salt<br>bridges |
| *Ca. L. solanacearum* | pyruvate | 7LOY | 1520 | 17 | 5 | 943 | 7 | 6 |
|  | pyruvate/SSA | 8GEK | 1485 | 18 | 5 | 912 | 5 | 6 |
|  | lysine | 7LVL | 1556 | 23 | 4 | 923 | 5 | 6 |
| *E. coli* | apo | 1YXC | 1289 | 9 | - | 497 | 6 | - |
|  | pyruvate | 3DU0 | 1226 | 9 | - | 478 | 4 | - |
|  | pyruvate/SSA | 4EOU | 1318 | 10 | - | 498 | 4 | - |
|  | lysine | 1YXD | 1297 | 9 | - | 488 | 7 | - |
| *A. tumefaciens* | apo | 4I7U | 1399 | 15 | 4 | 920 | 11 | 9 |
|  | pyruvate | 4I7V | 1394 | 15 | 4 | 950 | 14 | 15 |
| *B. anthracis* | apo | 1XL9 | 1393 | 10 | - | 872 | 7 | 4 |
|  | pyruvate | 3HIJ | 1392 | 10 | - | 872 | 7 | 4 |
| *T. maritima* | apo | 3PB2 | 1384 | 9 | - | 878 | 20 | 24 |

The weak dimer interface between monomers *a/c* (and *b/d*) includes residues Ile168, Glu169, Val197 and His198 that connect the two dimers (**Figure 5E**). Compared to the tight-dimer interface, the weak-dimer interface has on average six hydrogen bonds and six salt bridges. This is increased compared to *Ec*DHDPS, but consistent with other bacteria DHDPS enzymes (**Table 3**). The weak-dimer interface has been proposed to be a target for DHDPS inhibitor design (Mitsakos et al. 2008), since mutational studies of *E. coli* and *B. anthracis* DHDPS demonstrate that the formation of the tetramer is essential for enzyme activity (Griffin et al. 2008; Voss et al. 2010). Further, the weak-dimer interface is a region of low conservation across bacterial species. For *C*LsoDHDPS, the residues comprising the weak-dimer interface are similar to *A. tumefaciens,* but different to those in *E. coli*.

Although we were unable to crystalise the apo-*C*LsoDHDPS, we can overlay and compare the solution small-angle X-ray scattering data for *C*LsoDHDPS with and without various combinations of ligands to test for conformational changes upon ligand binding. Four data sets were collected and compared: apo-*C*LsoDHDPS, *C*LsoDHDPS + pyruvate, *C*LsoDHDPS + pyruvate/succinic semi-aldehyde, and *C*LsoDHDPS + lysine (**Figure 5F** and collection/analysis statistics in **Supplementary Table ii**). The scattering data were collected to a high S range (0.005–0.5 S), meaning we can reasonably expect to detect even small deviations in the global structure. Since the scattering plots overlay very closely, we conclude that *C*LsoDHDPS does not change in structure upon substrate binding, consistent with the crystal structures. That there was little change in *C*LsoDHDPS structure when the allosteric inhibitor lysine is bound, suggests that its mechanism of allosteric inhibition must require only subtle structural changes (if any at all), which is also found for *Ec*DHDPS (Blickling, Renner, et al. 1997; Blickling and Knäblein 1997; Dobson et al. 2005), but in contrast to the allosteric mechanism found in *N. sylvestris* DHDPS (Blickling, Beisel, et al. 1997). In the sections that follow, we will also demonstrate that the small-angle X-ray scattering data and the crystal structures match closely, evidence that the crystal structures faithfully represent the solution structures and are not affected by the crystal packing artifacts.

Overall, we demonstrate that the oligomeric state of *C*LsoDHDPS is tetrameric, as is the case for most other bacterial homologues. However, the interfaces between the monomers in *C*LsoDHDPS have increased surface area with more contacts, particularly across the functionally important tight-dimer interface. Further, we demonstrate that ligand binding does not alter the oligomeric structure in any appreciable way. This suggests that any destabilising effects of mutations accumulated as a result genetic drift is compensated for by increasing surface area and interactions at the tetrameric interfaces, a phenomenon suggested for enzymes that function at high temperatures (Vieille and Zeikus 2001; Reed et al. 2013).

### The CLsoDHDPS lysine and pyruvate binding sites are structurally the same as other DHDPS homologues

In an effort to explain the contrasting kinetic properties of *C*LsoDHDPS compared to homologues (*i.e.*, ternary-complex kinetic mechanism, slow turnover and high affinity for its substrates), we undertook a comparative analysis of residues that form the active and allosteric sites. We hypothesised that structural rearrangements in these sites, in particular, could account for these observations. A close examination of the *C*LsoDHDPS protein sequence, however, reveals that the residues within the allosteric and active site residues are largely retained (**Supplementary Figure ii**).

In plants and many bacteria, DHDPS catalyses the first committed step towards lysine biosynthesis (branchpoint reaction) and is feedback regulated by the allosteric binding of lysine, the pathway product. We have demonstrated that *C*LsoDHDPS is weakly inhibited by lysine (**Supplementary Figure iii**), similar to homologues from other Gram-negative bacteria. To determine how lysine binds to the allosteric cavity, which is ∼15 Å away from the active site, we solved the crystal structure with bound lysine to 2.01 Å resolution. The presence and orientation of two lysine molecules in each of the allosteric sites was confirmed from an omit map (**Supplementary Figure ivA**) and the refined (2F_o_-F_c_) density allowed accurate modelling of the binding pose (**Figure 6A**). Small angle X-ray scattering data confirms that the crystal structure faithfully represents the solutions structure (**Supplementary Figure ivB**). An overlay with other Gram-negative bacterial DHDPS homologues that are also weakly inhibited by lysine (*E. coli, A. tumefaciens*, and *C. jejuni*, **Figure 6B**) shows a highly conserved lysine binding pocket and the two lysine molecules engaging the site in an identical way, which is consistent with the similar low micromolar affinity for lysine. Although a definitive mechanism of allosteric inhibition has not been proposed for these homologues, it is likely that *C*LsoDHDPS uses an analogous mechanism.

**Figure 6.**
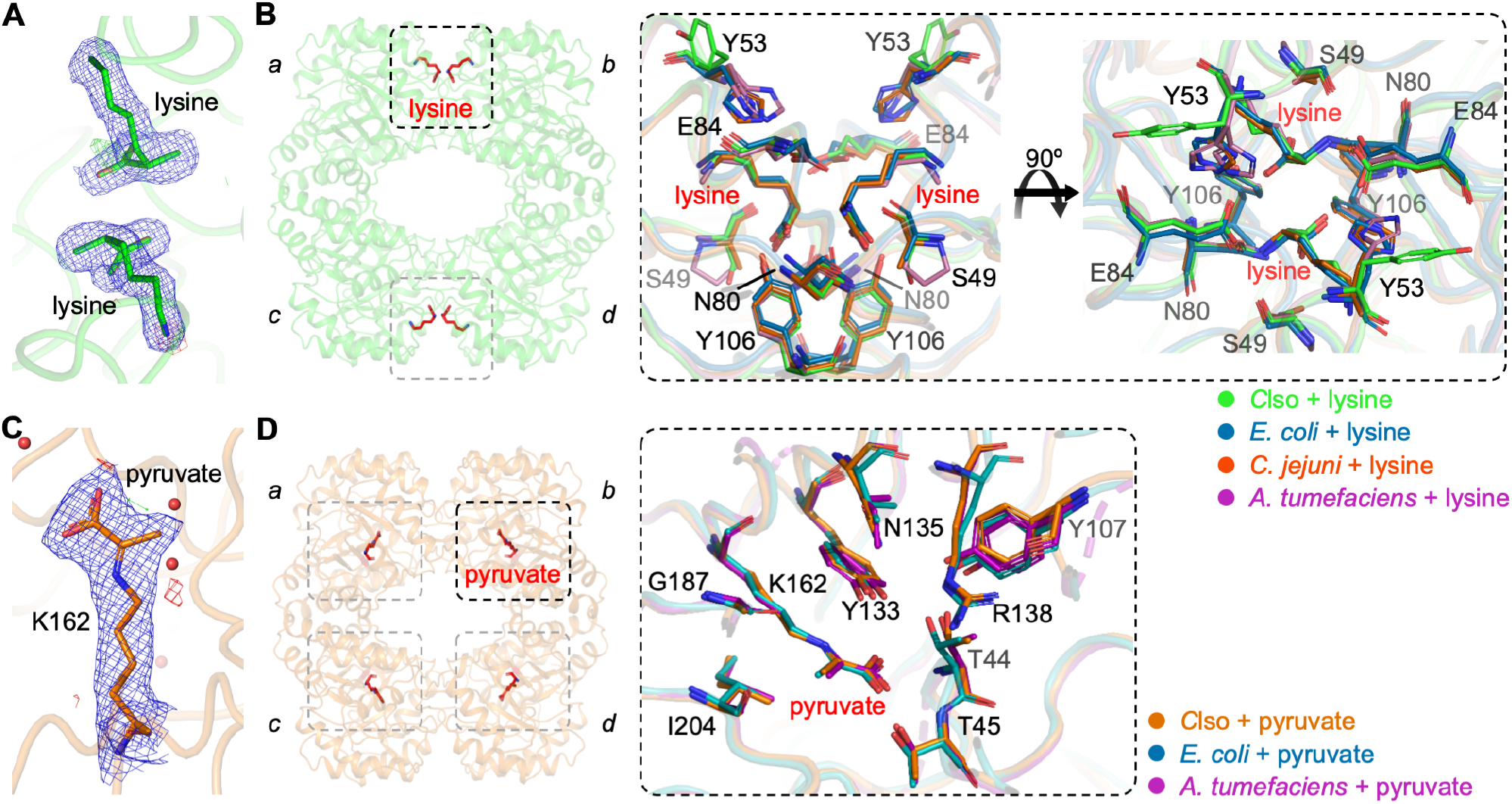
Structures of *C*LsoDHDPS with lysine or pyruvate. **A.** An electron density map [2F_o_-F_c_ at 1 σ (blue), F_o_-F_c_ at 3 σ (green) and −3 σ (red)] showing two lysine molecules within the cavity that forms the allosteric binding site. **B.** The *C*LsoDHDPS tetramer showing lysine (red sticks) bound in the allosteric site at the tight-binding interface formed through monomers *a*/*b* (or *c*/*d*). An enlargement of the allosteric site highlights residues that contact lysine, including Y53, E84, N80, Y106, which are within 4 Å. *C*LsoDHDPS with lysine (green, PDB ID: 7LVL) aligned with DHDPS homologues from *E. coli* (blue, PDB ID: 1YXD), *C. jejuni* (red, PDB ID: 4M19), and *A. tumefaciens* (magenta, PDB ID: 4I7W). In other weakly inhibited DHDPS homologues, Y53 is replaced with a histidine. **C.** Pyruvate covalently bound to K162 within the active site. Several water molecules are also present close to the K162-pyruvate adduct. **D.** The *C*LsoDHDPS tetramer showing pyruvate (red sticks) bound within the active site of each monomer. An enlargement and alignment of the catalytic site residues for *C*LsoDHDPS with pyruvate bound to K162 (orange, PDB ID: 7LVL) aligned with equivalent structures from *E. coli* (blue, PDB ID: 3DU0), and *A. tumefaciens* (magenta, PDB ID: 4I7V) demonstrate highly conserved contacts for this substrate.

Next, we solved the structure of *C*LsoDHDPS with the substrate pyruvate to 2.40 Å resolution. An omit map analysis (**Supplementary Figure ivC**) confirms that pyruvate is within the active site close to Lys162 and small angle X-ray scattering data confirms that the crystal structure faithfully represents the solutions structure (**Supplementary Figure ivD**). The refined structure reveals continuous (2F_o_-F_c_) electron density between Lys162 and pyruvate demonstrating that the enamine has formed (**Figure 6C**). The carboxylate group of pyruvate hydrogen bonds the side-chain hydroxyl of Thr45 and the mainchain nitrogen’s of Thr44 and Thr45 (**Figure 6D**). Pyruvate is positioned in a parallel plane to Tyr133 with the tyrosine hydroxyl group sitting above the enamine and Tyr133 is therefore well positioned to assist the formation of the Schiff base and subsequent enamine. The pyruvate methyl group points out from the active site, which places the enamine tautomer in an ideal position to attack the aldehyde of the incoming *S*-aspartate-β-semialdehyde substrate. An overlay with homologous structures with pyruvate reveals a highly similar active site architecture (**Figure 6D**) making it difficult to explain the decreased *K*_M_ for pyruvate (increased affinity). However, it is likely that the mechanism for the formation of the enamine is similar to other DHDPS enzymes. Lastly, we proposed above that the ternary-complex mechanism had ordered substrate binding [pyruvate binding followed by *S*-aspartate-β-semialdehyde binding]; that we find pyruvate bound to the active site in the enamine form is consisted with this assertion.

Together, these structures demonstrate that *C*LsoDHDPS retains the key residues for both allosteric inhibition by lysine and catalytic activity. Neither lysine nor pyruvate binding alter the structure in any appreciable way and the binding sites and binding poses of lysine and pyruvate are highly similar to other DHDPS homologues. From these data alone, it is difficult to rationalise the differences in function of *C*LsoDHDPS compared with other DHDPS homologues.

### CLsoDHDPS with bound pyruvate + succinic semi-aldehyde reveals a different product

We then determined the 1.93 Å resolution structure of *C*LsoDHDPS with bound pyruvate + succinic semi-aldehyde, which is an analogue of the second substrate *S*-aspartate-β- semialdehyde in that it lacks the primary amine. The rationale for this experiment is that since there are limited changes in the structure of *C*LsoDHDPS with pyruvate or lysine, perhaps the explanation for the unusual kinetic behaviours stems from the enzyme interaction with and catalysis of the second substrate *S*-aspartate-β-semialdehyde. Here, we expect the nucleophilic enamine (Schiff base) adduct of Lys162-pyruvate to attack the electrophilic carbon of the aldehyde of succinic semi-aldehyde to form a carbon-carbon covalent bond, as would occur with the natural substrate *S*-aspartate-β-semialdehyde—this occurs *in crystal* for the *Ec*DHDPS (Blickling, Renner, et al. 1997; Boughton et al. 2012). An omit map analysis confirms the presence of density consistent with pyruvate and succinic semi-aldehyde close to Lys162 (**Supplementary Figure ivE**) and again small angle X-ray scattering data confirms that the crystal structure faithfully represents the solutions structure (**Supplementary Figure ivF**).

Examination of the refined 2F_o_-F_c_ electron density map (**Figure 7A**) shows continuous density between Lys162, pyruvate and succinic semi-aldehyde such that the carbon-carbon bond between these substrates has formed. There is also clear density for the hydroxyl at the C4 position. The resolution of the data is sufficient to unambiguously model the hydroxyl in the *R*-configuration. In contrast, analogous structures of *Ec*DHDPS + pyruvate + succinic semi-aldehyde from separate research groups demonstrate that this hydroxyl is in the opposite *S*-configuration (Blickling, Renner, et al. 1997; Boughton et al. 2012). On comparing the refined 2F_o_-F_c_ electron density maps of *C*LsoDHDPS (**Figure 7A**) with pyruvate + succinic semi-aldehyde with the analogous structure of *Ec*DHDPS (**Figure 7B**), it is clear that the change in stereochemistry at this position results in the C4 hydroxyl sitting in a different position and pointing in a different direction. This is more obvious when we overlayed the active sites for both structures (**Figure 7C**). It is this difference that provides insight into the altered kinetics properties.

**Figure 7.**
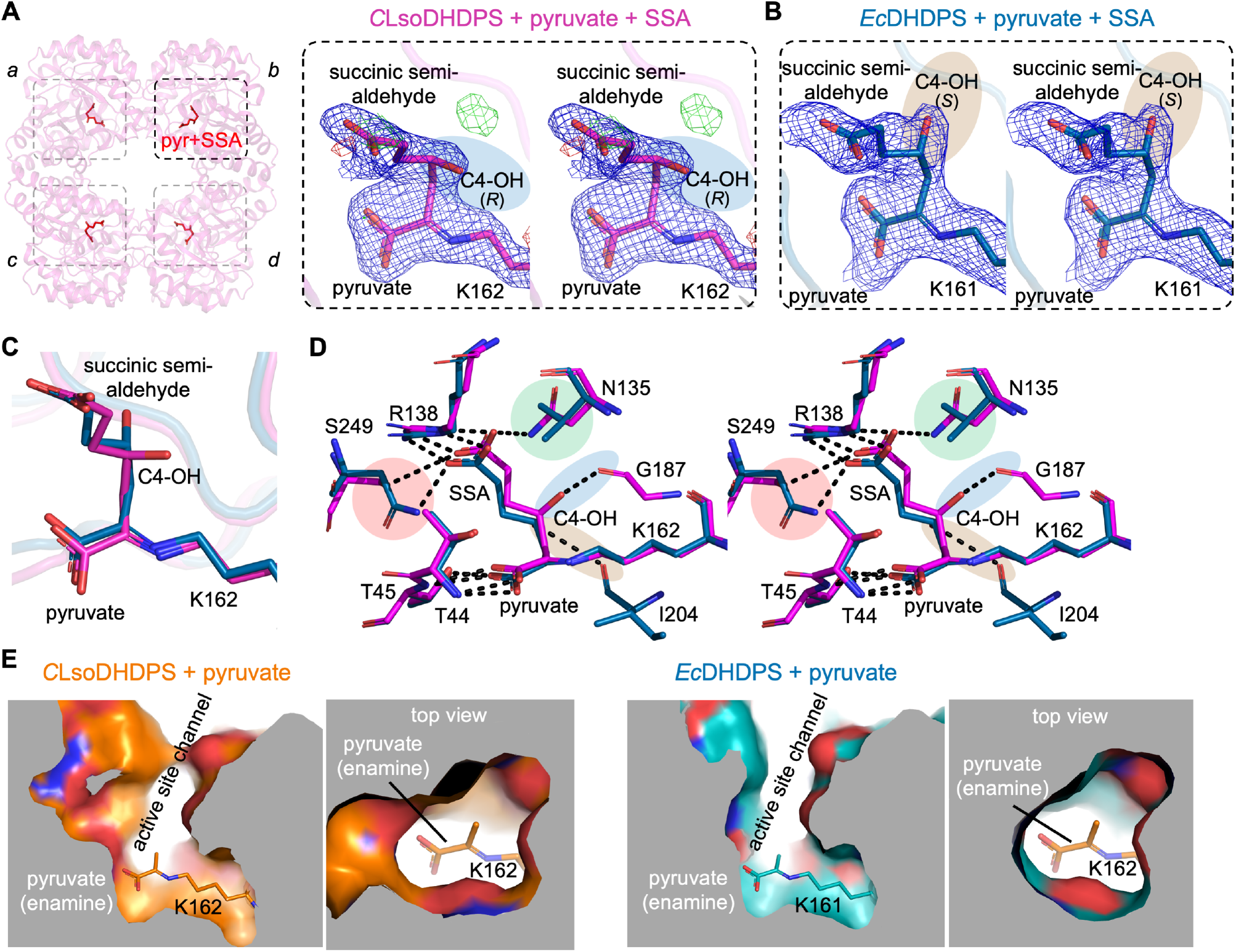
Structure of pyruvate + succinic semi-aldehyde bound *C*LsoDHDPS. **A.** and **B.** Cross-eyed stereo views comparing the refined electron density maps for the pyruvate + succinic semi-aldehyde bound *C*LsoDHDPS and *Ec*DHDPS active sites, demonstrating the different stereochemical configurations of the C4-hydroxyl (C4-OH, highlighted with blue shading in *C*LsoDHDPS and brown shading in *Ec*DHDPS). The σ values for the electron density maps are as follows: the 2F_o_-F_c_ map at 1 σ (blue) and the F_o_-F_c_ at 3 σ (green) and −3 σ (red). **C.** A focus on the covalently bound ligands (pyruvate and succinic semi-aldehyde) emphasising the difference in stereochemistry within the active site. Residue numbering as per *C*LsoDHDPS. **D.** An alignment and comparison of the active sites of *C*LsoDHDPS (magenta) and *Ec*DHDPS (blue, PDB ID:4EOU) with pyruvate + succinic semialdehyde adducts covalently bonded to the catalytic lysine in different stereochemical configurations. Residue numbering as per *C*LsoDHDPS. **E.** Cutaway views of the *C*LsoDHDPS and *Ec*DHDPS active sites with pyruvate, demonstrating the larger, more open volume that facilitates *S*-aspartate-β-semialdehyde binding in for *C*LsoDHDPS. The orientations for the *C*LsoDHDPS and *Ec*DHDPS active sites, relative to the pyruvate/lysine adduct, are identical, as is the slab size and position.

When comparing the interactions that the substrates make in the active site (**Figure 7D**) the hydroxyl group at the C4 position in the *Ec*DHDPS structure forms a polar contact with the main-chain of Ile203 (3.2 Å, highlighted with brown shading). In contrast, the altered stereochemistry of the C4-hydroxyl in *C*LsoDHDPS results in a close polar contact with the main chain carbonyl of Gly187 instead (∼2.76 Å, averaged over all monomers in the asymmetric unit, blue shading), and no connection to the equivalent Ile206 (∼4.6 Å distant, averaged over all monomers in the asymmetric unit). The difference in stereochemistry must be a result of the aldehyde from the second substrate *S*-aspartate-β-semialdehyde engaging the active site in a different conformation. That is, compared to the mechanism that operates in the *Ec*DHDPS homologue, the aldehyde is positioned such that nucleophilic attack must be from the opposite side of the carbonyl, resulting in the opposite stereochemical configuration. This has several mechanistic consequences that explain the altered kinetics and stereochemistry of the product.

1. Differences in residues that bind *S*-aspartate-β-semialdehyde are consistent with an increased affinity (decreased *K*_M_) for this substrate, particularly the carboxyl and amine moieties that position the aldehyde prior to condensation with pyruvate (**Supplementary Figure v**). For both *C*LsoDHDPS and *E*cDHDPS, the carboxyl of succinic acid interacts closely with Arg138 (O—N = ∼2.8 Å) and this residue is strictly conserved in DHDPS enzymes (**Supplementary Figure ii**). However, Asn135 and Ser249 in *C*LsoDHDPS are equivalent to Val135 and Asn248 in *E*cDHDPS (**Figure 7D**), which are also highly conserved residues across DHDPS enzymes, except for *Candidatus* Liberibacter species (**Supplementary Figure ii**). In *Ec*DHDPS the carboxyl of succinic acid contacts Asn248 (O—N = 2.7 Å), but in *C*LsoDHDPS it contacts both Ser249 (O—O = ∼4.2 Å) and Asn135 (O—N = 2.9 Å); this additional hydrogen bond is consistent with the increased affinity (decreased *K*_M_) for *S*-aspartate-β-semialdehyde.
2. The difference in succinic semialdehyde binding and larger active site explains the difference in the stereochemistry of the product. Because the pyruvate enamine structures are the same across all DHDPS homologues, including *C*LsoDHDPS (**Figure 6D**), it follows that *S*-aspartate-β-semialdehyde must bind the active site differently in order to present the aldehyde (electrophile) in a different orientation to provide the (*R,S*)-isomeric product. The alternate binding pattern places succinic acid in a slightly different position in the active site, presumably also placing the aldehyde in a different position prior to carbon-carbon bond formation, resulting in the different stereochemical outcome (**Supplementary Figure v**). Comparing the volume and shape of the active sites between *C*LsoDHDPS and *Ec*DHDPS (**Figure 7E**) reveals that the active site cavity for *C*LsoDHDPS is significantly wider at the *S*- aspartate-β-semialdehyde binding site, allowing *S*-aspartate-β-semialdehyde more conformational flexibility within the *C*LsoDHDPS active site, again consistent with the alternate stereochemical outcome of the product. A recent review concluded that active site linked tunnels and channels in enzymes affect activity, specificity, promiscuity, enantioselectivity and stability (Kokkonen et al. 2019). The wider active site channel of *C*LsoDHDPS could also allow faster and more efficient passage of substrates into and products out of the active site, consistent with the lower *K*_M_ values for the substrates.
3. There are two reasons why tighter *S*-aspartate-β-semialdehyde binding in the active site may lead to decreased *k*_cat_—a trade-off. Firstly, the condensation reaction between the pyruvate enamine and *S*-aspartate-β-semialdehyde is thought to be the rate determining step in the catalytic mechanism because it involves carbon-carbon bond formation. In *Ec*DHDPS, Ile203 (*Ec*DHDPS numbering) binds the hydroxyl in the pyruvate succinic semialdehyde adduct and is proposed to play a key role in this step (Dobson et al. 2008). In *C*LsoDHDPS, however, the C4-hydroxyl binds instead Gly186 and not Ile206 (*C*LsoDHDPS numbering) meaning that the isoleucine main chain does not participate during catalysis. Secondly, the final step of catalysis is the cyclisation through *S*-aspartate-β-semialdehyde amine. If the *S*-aspartate-β-semialdehyde carboxyl is constrained by additional bonding to Asn135 and Ser249, as seen in the *C*LsoDHDPS succinic acid bound structure, then the cyclisation step may be slowed.
4. Next, we considered the energetic consequences of the different products, which provides a surprising insight into the evolution of DHDPS enzymes more broadly. Quantum chemical modelling reveals that (4*S*)-hydroxy-2,3,4,5-tetrahydro-(2*S*)-dipicolinate forms an intramolecular hydrogen bond that preferentially stabilises this conformer by ∼12 kJ/mol compared to the (4*R*)-hydroxy-2,3,4,5-tetrahydro-(2*S*)-dipicolinate isomer produced by *C*LsoDHDPS-catalysed condensation of *S*-aspartate-β-semialdehyde and pyruvate (**Figure 8A**). A fundamental feature of enzymes is that they do not affect the equilibrium of the reaction (energy of the substrates versus the products), but instead catalyse the rate at which equilibrium is achieved (decrease the energy of the transition state) (Cornish-Bowden 1995; Fersht 1999). Since the product is both structurally and energetically different and therefore the equilibrium for this step has changed, evolution has in this case fashioned a new enzyme.
5. The intramolecular hydrogen bond present in the (*S,S*)-isomer, but not the (*R,S*)-isomer, also acts as a catalyst for the subsequent solution-phase dehydration reaction, providing a lower-energy pathway (decreased transition state energy) to the (*S*)-2,3-dihydropicolinate product (**Figure 8B**). This means that the subsequent dehydration step in the pathway is also changed, as it too will be slower compared to other bacterial diaminopimelate pathways, where the intramolecular hydrogen bond catalyses dehydration. Given that the product of the spontaneous dehydration is the same, the pathway likely continues to diaminopimelate and lysine as in other bacteria.

**Figure 8.**
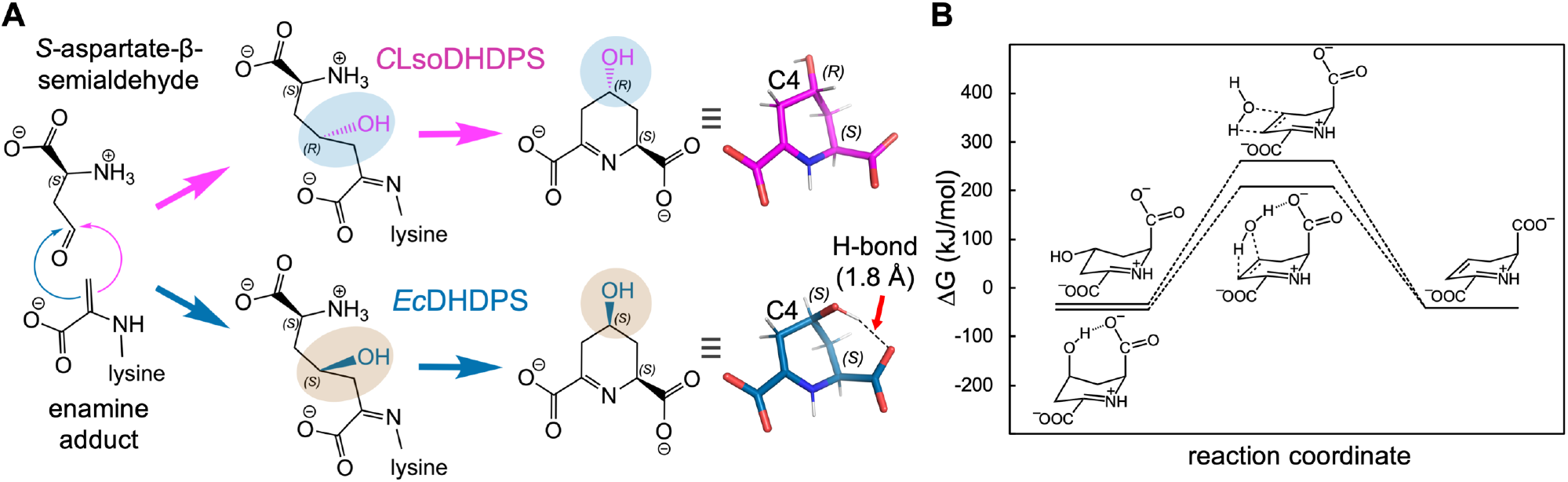
Reaction and energetics. **A.** The reaction pathways that generate the different stereochemical configurations of the product (4*R/S*)-hydroxy-2,3,4,5-tetrahydro-(2*S*)- dipicolinate. The C4-OH is coloured and shaded as in Figure 7A and B. To the right of the pane are the lowest energy conformers for each product, which reveals that only the (4*S*,2*S*)- configuration can form a hydrogen bond between the C4-OH and the C1-COO^-^, which preferentially stabilizes this isomer by ∼12 kJ/mol [based on computational benchmarking of analogous systems (Řezáč 2020)]. **B.** Reaction coordinate diagram for the solution-phase dehydration of (4*R*)-hydroxy-2,3,4,5-tetrahydro-(2*S*)-dipicolinate (*C*LsoDHDPS, top pathway) and (4*S*)-hydroxy-2,3,4,5-tetrahydro-(2*S*)-dipicolinate (*E*cDHDPS, bottom pathway) to form (*S*)-2,3-dihydropicolinate. All free energies are computed relative to the substrates of the DHDPS-catalysed reaction, (*S*)-aspartate-b-semialdehyde and pyruvate. The activation energy barrier (Δ_act_*G*°) for dehydration of (4*S*)-hydroxy-2,3,4,5-tetrahydro-(2*S*)-dipicolinate (*E*cDHDPS) is ∼130 kJ/mol, but for (4*R*)-hydroxy-2,3,4,5-tetrahydro-(2*S*)-dipicolinate (*C*LsoDHDPS) it is much higher at ∼175 kJ/mol.

To summarise, our high-resolution structure of *C*LsoDHDPS with bound pyruvate + succinic semi-aldehyde (1.93 Å) demonstrates an alternative stereochemistry for the product in the active site. This is driven by subtle changes in the *S*-aspartate-β-semialdehyde binding site that explains the increase affinity for the substrate, a lower turnover, and for a different stereochemical product. The (*R,S*)-isomer product of *C*LsoDHDPS is energetically less stable and has a higher energy for dehydration compared to the (*S,S*)-isomer product of other DHDPS enzymes, representing a fundamental change in the chemistry.

## Conclusions

Our objective was to define what effects genome reduction and population bottlenecks through vertical transmission may have on enzyme structure and function. Such studies are rare because proteins from reduced genomes tend to be highly unstable and therefore challenging to work with. As yet, there are no molecular structures for *Ca*. L. solanacearum proteins reported and structures from Liberibacter species in general are limited (Jiang et al. 2014; Saini et al. 2018; Kumar, Kesari, et al. 2019; Kumar, Dalal, et al. 2019; Nan et al. 2020), which hinders our understanding of the biology of this genus. This investigation to express, purify and characterise the ‘*Candidatus* Liberibacter solanacearum’ lysine biosynthetic enzyme dihydrodipicolinate synthase, which is well studied in other bacterial systems, provides a unique opportunity to assess how evolution shapes enzymes in context of a reduced genome and a bottlenecked lifestyle.

It is difficult to untangle the effects of natural selection (horizontal transmission *via* the plant) and drift (bottlenecking during successive vertical transmission) given the complex lifestyles of *Ca*. L. solanacearum. The genome characteristics are similar to bacteria such as *B. aphidicola* that have a high AT content and reduced size. Moreover, the aggregation propensity of *C*LsoDHDPS is certainly increased compared to homologues from bacteria with significantly larger genomes that are expected to evolve through natural selection. Indeed, this trend is also seen across other enzymes in the lysine biosynthetic pathway.

Our data supports the hypothesis that increased chaperone activity counters protein instability resulting from the accumulation of deleterious mutations. In addition, for *C*LsoDHDPS at least, instability is also overcome through increasing both the buried surface areas in oligomeric interfaces and the number of hydrogen bonds and salt bridges, an adaptation akin to that found in thermophilic enzymes that operate at high temperatures. Comparing *C*LsoDHDPS with the thermophilic homologue from *T. maritima*, demonstrates that the catalytically important tight-dimer interface, which comprises residues that form the active site and the allosteric binding site, is significantly stabilised, whereas in *Tm*DHDPS it is the weak-dimer interface that has increased stability. This perhaps reflects a need for *Tm*DHDPS when operating at high temperatures to retain its tetrameric state, which has been demonstrated to be important for function though minimising the protein dynamics through the dimer (and therefore the active sites). In contrast, *C*LsoDHDPS has stabilised the tight-binding interface directly as a way to overcome destabilising mutations that may occur due to drift.

Surprisingly, *C*LsoDHDPS is mechanistically the most divergent of all DHDPS enzymes studied to date. It binds its substrates with significantly higher affinity (low *K*_M_), yet is comparatively slow (low turnover); that is, there appears to be a trade-off between an increased binding affinity for the substrates and decreased catalysis. Moreover, given that *C*LsoDHDPS catalyses a reaction that results in a different product [(4*R*)-hydroxy-2,3,4,5- tetrahydro-(2*S*)-dipicolinate] that is energetically different (meaning the equilibrium for catalysis is different), it is in fact a new enzyme. Detailed structural data provides a molecular basis for these differences, whereby subtle differences in the active site affect binding and catalysis. We suggest that bacteria with endosymbiotic lifestyles present a rich source of interesting and variable enzymes useful for understanding enzyme function and/or informing protein engineering efforts.

The mechanistic differences of *C*LsoDHDPS also provides a new perspective to understand how DHDPS enzymes may have evolved in organisms expected to be under stringent natural selection. Specifically, these DHDPS enzymes likely evolved to afford the product (**4*S***)-hydroxy-2,3,4,5-tetrahydro-(2*S*)-dipicolinate, because it is more stable and able to catalyse the dehydration of the hydroxyl to form the substrate for the next enzyme in the pathway. In contrast, although the (**4*R***)*-*hydroxyl product is destabilised and cannot catalyse the dehydration in solution, the trade-off may be an active site that binds its substrate with higher affinity, which in the context of niches where *C*LsoDHDPS functions (plant phloem and insect gut) may be an advantage.

We see two future directions triggered by our work. Firstly, to study the effect on drift on enzymes requires a context where drift dominates. Our next work will focus on DHDPS enzymes from *Wolbachia pipientis*, *Buchnera aphidicola* and *Candidatus* Carsonella ruddii to unpick the effect of drift on these enzymes. Secondly, *Ca*. L. solanacearum has significant economic impact on the potato industries in Central and North America, and New Zealand. Novel disease management strategies will require an understanding of the structure and function of essential proteins required for pathogen survival in order to assist in the design of antimicrobials against this plant pathogen. Our kinetic and structural studies demonstrate that *C*LsoDHDPS is the most unusual of DHDPS enzyme studied to date. These differences, including kinetic mechanism, structure of the active site and substrate binding, can be exploited to develop specific inhibitors of *C*LsoDHDPS activity, but not of the corresponding enzyme in the plants they infect.

## Experimental

### Materials

The substrate *S*-aspartate-β-semialdehyde was synthesized using the previous methods (Roberts et al. 2003). The *E. coli* homologue of dihydrodipicolinate reductase was used in the coupled kinetic assay and was prepared using reported methods (Dobson, Valegård, et al. 2004).

### Gene expression analysis

#### Primer design

Primers were designed using the National Centre for Biotechnology Information (NCBI) Primer-Basic Local Alignment Search Tool (Primer-BLAST) (Ye et al. 2012). The genes and the primers used to amplify the gene products are described **Supplementary Tables iv** and **v**. Infected and uninfected psyllid and plant samples were tested for each primer pair in ensure primer specificity.

#### DNA extraction

Adult tomato potato psyllids were collected from *C*Lso-infected colonies. DNA was extracted from five psyllids from each colony using a cetyl trimethylammonium bromide extraction method (Beard and Scott 2013). DNA was also extracted from infected tomato plants using the cetyl trimethylammonium bromide extraction method (Beard et al. 2013). A semi-nested qPCR protocol (Beard and Scott 2013) was used in order to determine the titre of *Ca*. L. solanacearum present in DNA extracted from infected psyllids or tomato plants.

#### RNA extraction and cDNA synthesis

Gene expression experiments were performed using tomato plants, *Solanum lycopersicum* L. and psyllids infected with *Ca.* L. solanacearum. Prior to RNA extraction, plant samples were stored in RNA*later*™. The Geneaid Total RNA Mini Kit and the Aurum™ total RNA fatty and fibrous tissue kit were used for RNA extraction from psyllid and plant samples, respectively, as described by the manufacturer’s instructions. The isolated RNA was treated with RQ1 RNase-free DNase (Promega, Madison, WI) as described by the manufacturer. RNA was amplified using primers (described in **Supplementary Table iii**) by conventional PCR to test for the presence of any genomic DNA. cDNA was synthesized from 1 µg of total RNA using the iScript™ cDNA Synthesis Kit (BioRad) as described by the manufacturer.

#### Reverse transcription quantitative PCR (RT-qPCR)

All RT-qdPCR reactions were performed on a StepOne Plus™ real time PCR system (Thermo Fisher Scientific). SsoAdvanced™ universal SYBR reagent was mixed with appropriate primers at a final concentration of 0.3 µM and 5 µL cDNA in a reaction with a total volume of 20 µL. All samples were run in triplicate. Recombinase A (*recA,* KJZ80672) and the DNA-directed RNA polymerase beta subunit (*rpb,* KJZ81364.1) were used as reference genes to provide normalisation of relative quantification for the target gene.

#### Data analysis

The relative quantification of gene expression level was determined by the comparative C_T_ method 2^−ΔΔCT^ (Livak and Schmittgen 2001), where ^ΔΔ^C_T_ is calculated by the following equation:

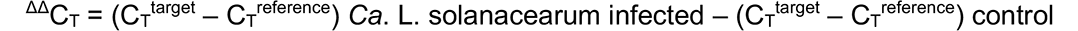

Standard deviation and P-values were calculated using a Student’s t-test (Yuan et al. 2006).

### Aggregation propensity

The AGGRESCAN algorithm (Conchillo-Solé et al. 2007) was used to predict the aggregation propensity of proteins in this study (http://bioinf.uab.es/aggrescan/). Amino acid sequences were submitted to AGGRESCAN in FASTA format and Na^4^vSS (Normalized a^4^v Sequence Sum for 100 residues) values were selected to compare the predictions.

### Protein expression and purification

The *C*LsoDHDPS was purified by the method previously described (Gilkes et al. 2019). Briefly, *E. coli* BL21 that harboured both the pGRO7 plasmid for GroEL and GroES expression as well as the pET24_*C*Lso_*dapA* plasmid, were grown overnight at 37 °C. Overnight cultures were used to inoculate 1 litre of Luria-Bertani broth and grown at 37 °C with shaking to an OD_600_ = 0.2, at which point chaperone expression was induced with L-arabinose at 0.5 mg/mL. The cultures were grown further to an OD_600_ of 0.5, at which point the protein of interest was induced with 1 mM IPTG and incubated at 16 °C for 20 hours with shaking. Protein concentration was measured by the Bradford method (Bradford 1976). Unless otherwise stated, enzymes were manipulated at 4 °C or on ice.

### Steady-state enzyme kinetics

DHDPS activity was measured using a dihydrodipicolinate reductase coupled assay, as previously described (Dobson, Valegård, et al. 2004). The reaction was initiated with *S*- aspartate-β-semialdehyde, after the cuvette had pre-incubated at 30 °C for ten minutes. Assays were performed in HEPES buffer (100 mM at pH 8) at 30 °C, kept constant using a circulating water bath. Care was taken to ensure that the enzymes and substrates were stable over the course of the assay. Care was also taken to ensure an excess of DHDPR; about 20 μg per assay was used. The *K*_M_ values for *C*LsoDHDPS were unknown; therefore, apparent *K*_M_ values were initially determined using a range of substrate concentrations. True kinetic constants (*K*_M_ and *k*_cat_) were determined using a range of concentrations approximately 10-fold above and below these values. Initial velocities were reproducible within 10%. All data were analysed using R. Data were fitted to the appropriate models as judged by the R^2^ value and the Akaike information criterion (AIC).

### Small angle X-ray scattering

Small angle X-ray scattering (SAXS) data were collected on the SAXS/WAXS beamline equipped with a Pilatus 1M detector (170 mm × 170 mm, effective pixel size, 172 μm × 172 μm) at the Australian Synchrotron. A sample detector distance of 1600 mm was used, providing a *q* range of 0.006–0.5 Å^-1^. The purified protein was injected onto an inline Superdex S200 Increase 5/150 GL SEC column (GE Healthcare), equilibrated with 1× PBS buffer (pH 7.4) supplemented with 0.1% (w/v) sodium azide, at a flow rate of 0.35 mL/min^-1^. Co-flow SAXS was used to minimise sample dilution and maximise signal to noise ratio. Scattering data were collected in 1 s exposures (λ = 1.0332 Å) over 500 frames, using a 1.5 mm glass capillary, at 12 °C.

Analysis of the scattering data were performed using the ATSAS software package (version 2.8.4) (Franke et al. 2017). The molecular mass of the samples was estimated using the SAXS-MoW2 package. 2D intensity plots were radially averaged, normalised to sample transmission, and background subtracted using CHROMIXS. Guinier plots were analysed using PRIMUS (Franke et al. 2017) to assess data quality. Indirect Fourier transform of the data were performed using GNOM (Svergun 1992) to generate the *P(r)* distribution. Theoretical scattering curves were generated from atomic coordinates and compared with experimental scattering curves using CRYSOL (Svergun et al. 1995). Data collection and analysis statistics are summarised in **Supplementary Table ii**.

### X-ray crystallography

Crystallisation studies were performed using freshly purified protein, concentrated to 14 mg/mL. Crystallisation trials used the sitting drop vapour diffusion method, at 20 °C, with droplets consisting of 150 nL of protein solution and 150 nL of reservoir solution. Rectangular *C*LsoDHDPS crystals were produced using the JCSG+ Suite screen in condition A2 (20% *w/v* polyethylene glycol 3000, 0.1 M sodium citrate, pH 5.5) at 20 °C. For ligand bound structures, the crystals were soaked for 7 days in mother liquor with either: 100 mM lysine, 100 mM pyruvate or 100 mM of both succinic semi-aldehyde and pyruvate. Prior to X-ray data collection, the crystals were soaked in cryoprotectant solution consisting of 85% reservoir solution and 15% glycerol-ethylene glycol (50:50 mix) prior to being flash frozen in liquid nitrogen.

Diffraction data were collected at 110 K on the MX2 beamline at the Australian Synchrotron (McPhillips et al. 2002) with the EIGER × 16M detector (λ = 0.9537 Å). All liganded crystals belong to the space group C2 with diffraction to 2.40 Å for *C*LsoDHDPS + pyruvate), 1.93 Å for *C*LsoDHDPS + pyruvate/SSA, and 2.01 Å for *C*LsoDHDPS + lysine (**Supplementary Table iii**). Diffraction data sets were processed and scaled using XDS and AIMLESS from the CCP4i2 program suite (Potterton et al. 2018). The resulting intensity data were analysed using PHENIX XTRIAGE (Adams et al. 2010). Molecular replacement was performed using PHASER (McCoy et al. 2007) with *A. tumefaciens* DHDPS (PDB ID 4I7U) as the search model. CHAINSAW (Stein 2008) from the CCP4i2 suite was used to prepare the model of *C*LsoDHDPS, omitting waters and reducing it to its monomeric form. Structural refinement of the resulting six monomers was performed using REFMAC5 (Murshudov et al. 1997) with iterative model building using COOT (Emsley and Cowtan 2004). Initial refinement initially applied non-crystallographic restraints followed by simulated annealing using PHENIX with the structure validated using MolProbity (Chen et al. 2010). Images were generated in PyMOL (DeLano 2002) or Chimera (Pettersen et al. 2021). Data collection and model-refinement statistics are in **Supplementary Table iii**.

### Computational modelling

Geometries of (4*S*)-hydroxy-2,3,4,5-tetrahydro-(2*S*)-dipicolinate, (4*R*)-hydroxy-2,3,4,5- tetrahydro-(2*S*)-dipicolinate, *S*-aspartate-β-semialdehyde, pyruvate and *S*-2,3- dihydropicolinate were optimised at B3LYP/6-31G*, considering all possible rotamers. The lowest energy conformers identified were confirmed as minima by the absence of imaginary frequencies following harmonic frequency calculations, from which zero-point (*H*_ZVPE_), enthalpic (*H*_therm_) and entropic (*S*_therm_) contributions to the free energy of each species under standard thermodynamic conditions were also obtained. Single point calculations were performed at ωB97X-V/def2-QZVPPD to more accurately estimate the electronic energy (*U*_elec_) and absolute free energies computed as:

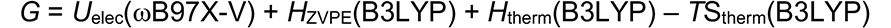

Free energies of reaction, ΔG, were computed relative to the free energy of (*S*)-aspartate-β- semialdehyde and pyruvate in solution. Transition states for eliminative dehydration were identified by performing 2D relaxed scans along the C-O and O-H bond length coordinates (step size = 0.2 Å), followed by an automated transition state optimisation. Frequency calculations were performed to confirm that these optimisations had converged a first-order transition state, indicated by the presence of a single imaginary frequency. Transition state free energies were computed as above, excluding the imaginary frequency vibrational mode that corresponds to the reaction coordinate. All calculations were performed under the influence of a polarisable continuum solvation model with a dielectric constant chosen to resemble water (ε = 78.36), using the QChem5.3 programme package (Epifanovsky et al. 2021).

## Acknowledgements

This research was undertaken in part using the MX2 beamline at the Australian Synchrotron, part of ANSTO, and made use of the Australian Cancer Research Foundation (ACRF) detector as well as the SAXS beamline at the Australian Synchrotron, part of ANSTO. J.M.G. is the recipient of a Callaghan Innovation PhD scholarship. R.C.J.D. acknowledges the following for funding support, in part: 1) the Marsden Fund council from Government funding, managed by Royal Society Te Apārangi (contract UOC1506); 2) a Ministry of Business, Innovation and Employment Smart Ideas grant (contract UOCX1706); and 3) the Biomolecular Interactions Centre (UC). R.A.F and G.R.S acknowledge the support of Plant & Food Research to undertake Liberibacter research via MBIE C11X1308, CAP10-72 and PBCRC2002/2156 funding.

## Data Availability Statement

The data underpinning this article are available in the article and in its online supplementary material. Any further data required associated with this article will be shared on reasonable request to the corresponding author.

## Notes

### Competing Interest Statement

The authors have declared no competing interest.

## References

Adams PD, Afonine PV, Bunkóczi G, Chen VB, Davis IW, Echols N, Headd JJ, Hung L-W, Kapral GJ, Grosse-Kunstleve RW, et al. 2010. PHENIX: a comprehensive Python-based system for macromolecular structure solution. Acta Crystallogr Sect D Biological Crystallogr 66:213–221.

Ammar E, Shatters RG, Hall DG. 2011. Localization of Candidatus Liberibacter asiaticus, associated with citrus huanglongbing disease, in its psyllid vector using fluorescence in situ hybridization. J Phytopathol 159:726–734.

Atkinson SC, Dogovski C, Downton MT, Pearce FG, Reboul CF, Buckle AM, Gerrard JA, Dobson RCJ, Wagner J, Perugini MA. 2012. Crystal, solution and in silico structural studies of dihydrodipicolinate synthase from the common grapevine. PLoS One 7:e38318–9.

Atkinson SC, Dogovski C, Wood K, Griffin MDW, Gorman MA, Hor L, Reboul CF, Buckle AM, Wuttke J, Parker MW, et al. 2018. Substrate locking promotes dimer-dimer docking of an enzyme antibiotic target. Structure 26:948–959.

Beard SS, Pitman AR, Kraberger S, Scott IAW. 2013. SYBR Green real-time quantitative PCR for the specific detection and quantification of ‘Candidatus Liberibacter solanacearum’ in field samples from New Zealand. Eur J Plant Pathol 136:203–215.

Beard SS, Scott IAW. 2013. A rapid method for the detection and quantification of the vector-borne bacterium ‘Candidatus Liberibacter solanacearum’ in the tomato potato psyllid, Bactericera cockerelli. Entomol Exp Appl 147:196–200.

Blickling S, Beisel H-G, Bozic D, Knäblein J, Laber B, Huber R. 1997. Structure of dihydrodipicolinate synthase of Nicotiana sylvestris reveals novel quaternary structure. J Mol Biol 274:608–621.

Blickling S, Knäblein J. 1997. Feedback inhibition of dihydrodipicolinate synthase enzymes by L-lysine. J Biol Chem 378:207–210.

Blickling S, Renner C, Laber B, Pohlenz H-D, Holak TA, Huber R. 1997. Reaction mechanism of Escherichia coli dihydrodipicolinate synthase investigated by X-ray crystallography and NMR spectroscopy. Biochemistry 36:24–33.

Boughton BA, Dobson RCJ, Hutton CA. 2012. The crystal structure of dihydrodipicolinate synthase from Escherichia coli with bound pyruvate and succinic acid semialdehyde: Unambiguous resolution of the stereochemistry of the condensation product. Proteins Struct Funct Bioinform 80:2117–2122.

Boughton BA, Griffin MDW, O’Donnell PA, Dobson RCJ, Perugini MA, Gerrard JA, Hutton CA. 2008. Irreversible inhibition of dihydrodipicolinate synthase by 4-oxo-heptenedioic acid analogues. Bioorg Med Chem 16:9975–9983.

Bradford MM. 1976. A rapid and sensitive method for the quantitation of microgram quantities of protein utilizing the principle of protein-dye binding. Anal Biochem 72:248– 254.

Buchman JL, Fisher TW, Sengoda VG, Munyaneza JE. 2012. Zebra chip progression: From inoculation of potato plants with Liberibacter to development of disease symptoms in tubers. Am J Potato Res 89:159–168.

Buchman JL, Sengoda VG, Munyaneza JE. 2011. Vector transmission efficiency of Liberibacter by Bactericera cockerelli (Hemiptera: Triozidae) in zebra chip potato disease: Effects of psyllid life stage and inoculation access period. J Econ Entomol 104:1486– 1495.

Burgess BR, Dobson RCJ, Bailey MF, Atkinson SC, Griffin MDW, Jameson GB, Parker MW, Gerrard JA, Perugini MA. 2008. Structure and evolution of a novel dimeric enzyme from a clinically important bacterial pathogen. J Biol Chem 283:27598–27603.

Casteel CL, Hansen AK, Walling LL, Paine TD. 2012. Manipulation of plant defense responses by the tomato psyllid (Bactericerca cockerelli) and its associated endosymbiont Candidatus Liberibacter psyllaurous. PLoS One 7:e35191.

Chen VB, Arendall WB, Headd JJ, Keedy DA, Immormino RM, Kapral GJ, Murray LW, Richardson JS, Richardson DC. 2010. MolProbity: all-atom structure validation for macromolecular crystallography. Acta Crystallogr Sect D Biological Crystallogr 66:12–21.

Chowdhury R, Sahu GK, Das J. 1996. Stress response in pathogenic bacteria. J Biosciences 21:149–160.

Christoff RM, Gardhi CK, Costa TPS da, Perugini MA, Abbott BM. 2019. Pursuing DHDPS: an enzyme of unrealised potential as a novel antibacterial target. MedChemComm 10:1581–1588.

Cicero JM, Fisher TW, Qureshi JA, Stansly PA, Brown JK. 2017. Colonization and intrusive invasion of potato psyllid by ‘ Candidatus Liberibacter solanacearum.’ Phytopathology 107:36–49.

Conchillo-Solé O, Groot NS de, Avilés FX, Vendrell J, Daura X, Ventura S. 2007. AGGRESCAN: a server for the prediction and evaluation of “hot spots” of aggregation in polypeptides. BMC Bioinform 8:65.

Cornish-Bowden A. 1995. Fundamentals of enzyme kinetics. Brookfield: Portland

Costa TPS da, Hall CJ, Panjikar S, Wyllie JA, Christoff RM, Bayat S, Hulett MD, Abbott BM, Gendall AR, Perugini MA. 2021. Towards novel herbicide modes of action by inhibiting lysine biosynthesis in plants. eLife 10:e69444.

Cox RJ. 1996. The DAP pathway to lysine as a target for antimicrobial agents. Nat Prod Rep, 13:29–43.

Cox RJ, Sutherland A, Vederas JC. 2000. Bacterial diaminopimelate metabolism as a target for antibiotic design. Bioorg Med Chem 8:843–871.

DeLano WL. 2002. Pymol: An open-source molecular graphics tool. CCP4 Newsl protein Crystallogr. 40:82–92.

Dobson RCJ, Griffin MDW, Devenish SRA, Pearce FG, Hutton CA, Gerrard JA, Jameson GB, Perugini MA. 2008. Conserved main-chain peptide distortions: a proposed role for Ile203 in catalysis by dihydrodipicolinate synthase. Protein Science 17:2080–2090.

Dobson RCJ, Griffin MDW, Jameson GB, Gerrard JA. 2005. The crystal structures of native and (S)-lysine-bound dihydrodipicolinate synthase from Escherichia coli with improved resolution show new features of biological significance. Acta Crystallogr Sect D Biological Crystallogr 61:1116–1124.

Dobson RCJ, Griffin MDW, Roberts SJ, Gerrard JA. 2004. Dihydrodipicolinate synthase (DHDPS) from Escherichia coli displays partial mixed inhibition with respect to its first substrate, pyruvate. Biochimie 86:311–315.

Dobson RCJ, Valegård K, Gerrard JA. 2004. The crystal structure of three site-directed mutants of Escherichia coli dihydrodipicolinate synthase: Further evidence for a catalytic triad. J Mol Bio 338:329–339.

Domigan LJ, Scally SW, Fogg MJ, Hutton CA, Perugini MA, Dobson RCJ, Muscroft-Taylor AC, Gerrard JA, Devenish SRA. 2009. Characterisation of dihydrodipicolinate synthase (DHDPS) from Bacillus anthracis. Biochim Biophys Acta Proteins Proteom 1794:1510– 1516.

Douglas AE. 2006. Phloem-sap feeding by animals: problems and solutions. J Exp Bot 57:747–754.

Emsley P, Cowtan K. 2004. Coot: model-building tools for molecular graphics. Acta Crystallogr Sect D Biological Crystallogr 60:2126–2132.

Epifanovsky E, Gilbert ATB, Feng X, Lee J, Mao Y, Mardirossian N, Pokhilko P, White AF, Coons MP, Dempwolff AL, et al. 2021. Software for the frontiers of quantum chemistry: An overview of developments in the Q-Chem 5 package. J Chem Phys 155:084801.

Fares MA, Barrio E, Sabater-Muñoz B, Moya A. 2002. The evolution of the heat-shock protein GroEL from Buchnera, the primary endosymbiont of aphids, is governed by positive selection. Mol Biol Evol 19:1162–1170.

Fares MA, Moya A, Barrio E. 2004. GroEL and the maintenance of bacterial endosymbiosis. Trends Genet 20:413–416.

Fares MA, Moya A, Barrio E. 2005. Adaptive evolution in GroEL from distantly related endosymbiotic bacteria of insects. J Evol Biol 18:651–660.

Fares MA, Ruiz-González MX, Moya A, Elena SF, Barrio E. 2002. GroEL buffers against deleterious mutations. Nature 417:398–398.

Fersht A. 1999. Structure and Mechanism in Protein Science: a Guide to Enzyme Catalysis and Protein Folding. New York: Freeman

Franke D, Petoukhov MV, Konarev PV, Panjkovich A, Tuukkanen A, Mertens HDT, Kikhney AG, Hajizadeh NR, Franklin JM, Jeffries CM, et al. 2017. ATSAS 2.8: a comprehensive data analysis suite for small-angle scattering from macromolecular solutions. J Appl Crystallogr 50:1212–1225.

Georgescauld F, Popova K, Gupta AJ, Bracher A, Engen JR, Hayer-Hartl M, Hartl FU. 2014. GroEL/ES chaperonin modulates the mechanism and accelerates the rate of TIM-barrel domain folding. Cell 157:922–934.

Gilkes JM, Sheen CR, Frampton RA, Smith GR, Dobson RCJ. 2019. The first purification of functional proteins from the unculturable, genome-reduced, bottlenecked α- proteobacterium ‘Candidatus Liberibacter solanacearum.’ Phytopathology 109:1141– 1148.

Girodeau JM, Agouridas C, Masson M, Pineau R, Goffic FL. 1986. The lysine pathway as a target for a new genera of synthetic antibacterial antibiotics? J Med Chem 29:1023–1030.

Griffin MDW, Dobson RCJ, Pearce FG, Antonio L, Whitten AE, Liew CK, Mackay JP, Trewhella J, Jameson GB, Perugini MA, et al. 2008. Evolution of quaternary structure in a homotetrameric enzyme. J Mol Bio 380:691–703.

Ham RCHJ van,Kamerbeek J, Palacios C, Rausell C, Abascal F, Bastolla U, Fernández JM, Jiménez L, Postigo M, Silva FJ, et al. 2003. Reductive genome evolution in Buchnera aphidicola. Proc National Acad Sci U.S.A. 100:581–586.

Hansen AK, Trumble JT, Stouthamer R, Paine TD. 2008. A new huanglongbing species, “Candidatus Liberibacter psyllaurous”, found to infect tomato and potato, Is vectored by the psyllid Bactericera cockerelli (Sulc). Appl Environ Microb 74:5862–5865.

Hutton CA, Perugini MA, Gerrard JA. 2007. Inhibition of lysine biosynthesis: an evolving antibiotic strategy. Mol Biosyst 3:458–465.

Ibanez F, Levy J, Tamborindeguy C. 2014. Transcriptome analysis of “Candidatus Liberibacter solanacearum” in its psyllid vector, Bactericera cockerelli. PLoS One 9:e100955.

Jiang L, Gao Z, Li Y, Wang S, Dong Y. 2014. Crystal structures and kinetic properties of enoyl-acyl carrier protein reductase I from Candidatus Liberibacter asiaticus. Protein Sci 23:366–377.

Kefala G, Evans GL, Griffin MDW, Devenish SRA, Pearce FG, Perugini MA, Gerrard JA, Weiss MS, Dobson RCJ. 2008. Crystal structure and kinetic study of dihydrodipicolinate synthase from Mycobacterium tuberculosis. Biochem J 411:351–10.

Kelkar YD, Ochman H. 2013. Genome reduction promotes increase in protein functional complexity in bacteria. Genetics 193:303–307.

Kelley AJ, Pelz-Stelinski KS. 2019. Maternal contribution of Candidatus Liberibacter asiaticus to asian citrus psyllid (Hemiptera: Liviidae) nymphs through oviposition site inoculation and transovarial transmission. J Econ Entomol 112:2565–2568.

Kokkonen P, Bednar D, Pinto G, Prokop Z, Damborsky J. 2019. Engineering enzyme access tunnels. Biotechnol Adv 37:107386.

Krissinel E, Henrick K. 2007. Inference of macromolecular assemblies from crystalline state. J Mol Bio 372:774–797.

Kumar P., Dalal V, Sharma N, Kokane S, Ghosh DK, Kumar Pravindra, Sharma AK. 2019. Characterization of the heavy metal binding properties of periplasmic metal uptake protein CLas-ZnuA2. Metallomics 12:280–289.

Kumar P., Kesari P, Kokane S, Ghosh DK, Kumar Pravindra, Sharma AK. 2019. Crystal structures of a putative periplasmic cystine-binding protein from Candidatus Liberibacter asiaticus: Insights into an adapted mechanism of ligand binding. FEBS J 286:3450–3472.

Lin H, Lou B, Glynn JM, Doddapaneni H, Civerolo EL, Chen C, Duan Y, Zhou L, Vahling CM. 2011. The complete genome sequence of ‘Candidatus Liberibacter solanacearum’, the bacterium associated with potato zebra chip disease. PLoS One 6:e19135.

Livak KJ, Schmittgen TD. 2001. Analysis of relative gene expression data using real-time quantitative PCR and the 2−ΔΔC T method. Methods 25:402–408.

Marais GAB, Calteau A, Tenaillon O. 2008. Mutation rate and genome reduction in endosymbiotic and free-living bacteria. Genetica 134:205–210.

McCoy AJ, Grosse-Kunstleve RW, Adams PD, Winn MD, Storoni LC, Read RJ. 2007. Phaser crystallographic software. J Appl Crystallogr 40:658–674.

McCutcheon JP, Moran NA. 2012. Extreme genome reduction in symbiotic bacteria. Nat Rev Microbiol 10:13.

McLennan N, Masters M. 1998. GroE is vital for cell-wall synthesis. Nature 392:139–139.

McPhillips TM, McPhillips SE, Chiu H-J, Cohen AE, Deacon AM, Ellis PJ, Garman E, Gonzalez A, Sauter NK, Phizackerley RP, et al. 2002. Blu-Ice and the distributed control system: Software for data acquisition and instrument control at macromolecular crystallography beamlines. J Synchrotron Radiat 9:401–406.

Mitsakos V, Dobson RCJ, Pearce FG, Devenish SR, Evans GL, Burgess BR, Perugini MA, Gerrard JA, Hutton CA. 2008. Inhibiting dihydrodipicolinate synthase across species: towards specificity for pathogens? Bioorg Med Chem Lett 18:842–844.

Moran NA. 1996. Accelerated evolution and Muller’s rachet in endosymbiotic bacteria. Proc. Natl. Acad. Sci. U.S.A. 93:2873–2878.

Munyaneza JE. 2010. Psyllids as vectors of emerging bacterial diseases of annual crops. Southwest Entomol 35:471–477.

Munyaneza JE. 2012. Zebra chip disease of potato: Biology, epidemiology, and management. Am J Potato Res 89:329–350.

Munyaneza JE, Crosslin JM, Upton JE. 2007. Further evidence that zebra chip potato disease in the lower Rio Grande valley of Texas is associated with Bactericera cockerelli. Subtrop plant Sci. 59:30–37.

Murshudov GN, Vagin AA, Dodson EJ. 1997. Refinement of macromolecular structures by the maximum-likelihood method. Acta Crystallogr Sect D Biological Crystallogr 53:240– 255.

Nachappa P, Levy J, Tamborindeguy C. 2012. Transcriptome analyses of Bactericera cockerelli adults in response to “Candidatus Liberibacter solanacearum” infection. Mol Genet Genomics 287:803–817.

Nakabachi A, Yamashita A, Toh H, Ishikawa H, Dunbar HE, Moran NA, Hattori M. 2006. The 160-kilobase genome of the bacterial endosymbiont Carsonella. Science 314:267–267.

Nan J, Zhang S, Zhan P, Jiang L. 2020. Evaluation of bronopol and disulfiram as potential candidatus Liberibacter asiaticus inosine 5′-monophosphate dehydrogenase inhibitors by using molecular docking and enzyme kinetic. Molecules 25:2313.

Pearce FG, Dobson RCJ, Jameson GB, Perugini MA, Gerrard JA. 2011. Characterization of monomeric dihydrodipicolinate synthase variant reveals the importance of substrate binding in optimizing oligomerization. Biochim Biophys Acta Proteins Proteom 1814:1900– 1909.

Pelz-Stelinski KS, Brlansky RH, Ebert TA, Rogers ME. 2010. Transmission parameters for Candidatus Liberibacter asiaticus by Asian citrus psyllid (Hemiptera: Psyllidae). J Econ Entomol 103:1531–1541.

Pettersen EF, Goddard TD, Huang CC, Meng EC, Couch GS, Croll TI, Morris JH, Ferrin TE. 2021. UCSF ChimeraX: Structure visualization for researchers, educators, and developers. Protein Sci 30:70–82.

Pitman AR, Drayton GM, Kraberger SJ, Genet RA, Scott IAW. 2011. Tuber transmission of ‘Candidatus Liberibacter solanacearum’ and its association with zebra chip on potato in New Zealand. Eur J Plant Pathol 129:389–398.

Potterton L, Agirre J, Ballard C, Cowtan K, Dodson E, Evans PR, Jenkins HT, Keegan R, Krissinel E, Stevenson K, et al. 2018. CCP4i2: the new graphical user interface to the CCP4 program suite. Acta Crystallogr Sect D 74:68–84.

Reed CJ, Lewis H, Trejo E, Winston V, Evilia C. 2013. Protein Adaptations in Archaeal Extremophiles. Archaea 2013:373275.

Řezáč J. 2020. Non-covalent interactions atlas benchmark data sets: Hydrogen bonding. J Chem Theory Comput 16:2355–2368.

Rice EA, Bannon GA, Glenn KC, Jeong SS, Sturman EJ, Rydel TJ. 2008. Characterization and crystal structure of lysine insensitive Corynebacterium glutamicum dihydrodipicolinate synthase (cDHDPS) protein. Arch Biochem Biophys 480:111–121.

Roberts SJ, Morris JC, Dobson RCJ, Gerrard JA. 2003. The preparation of (S)-aspartate semi-aldehyde appropriate for use in biochemical studies. Bioorg Med Chem 13:265–267.

Rutherford SL. 2003. Between genotype and phenotype: protein chaperones and evolvability. Nat Rev Genet 4:263–274.

Saini G, Sharma N, Dalal V, Warghane A, Ghosh DK, Kumar P, Sharma AK. 2018. The analysis of subtle internal communications through mutation studies in periplasmic metal uptake protein CLas-ZnuA2. J Struct Biol 204:228–239.

Secor GA, Lee I-M, Bottner KD, Rivera-Varas V, Gudmestad NC. 2006. First report of a defect of processing potatoes in Texas and Nebraska associated with a new phytoplasma. Plant Dis 90:377–377.

Stein N. 2008. CHAINSAW: a program for mutating pdb files used as templates in molecular replacement. J Appl Crystallogr 41:641–643.

Svergun D, Barberato C, Koch M. 1995. CRYSOL – a program to evaluate X-ray solution scattering of biological macromolecules from atomic coordinates. J Appl Crystallogr 28:768–773.

Svergun DI. 1992. Determination of the regularization parameter in indirect-transform methods using perceptual criteria. J Appl Crystallogr 25:495–503.

Thao ML, Moran NA, Abbot P, Brennan EB, Burckhardt DH, Baumann P. 2000. Co-speciation of psyllids and their primary prokaryotic endosymbionts. Appl Environ Microb 66:2898–2905.

Tokuriki N, Tawfik Dan S. 2009. Chaperonin overexpression promotes genetic variation and enzyme evolution. Nature 459:668–673.

Tokuriki N, Tawfik Dan S. 2009. Stability effects of mutations and protein evolvability. Curr Opin Struc Biol 19:596–604.

Turner JJ, Healy JP, Dobson RCJ, Gerrard JA, Hutton CA. 2005. Two new irreversible inhibitors of dihydrodipicolinate synthase: diethyl (E,E)-4-oxo-2,5-heptadienedioate and diethyl (E)-4-oxo-2-heptenedioate. Bioorg Med Chem Lett 15:995–998.

Vereijssen J, Smith GR, Weintraub PG. 2018. Bactericera cockerelli (Hemiptera: Triozidae) and Candidatus Liberibacter solanacearum in Potatoes in New Zealand: Biology, Transmission, and Implications for Management. J Integr Pest Manag 9:13.

Vieille C, Zeikus GJ. 2001. Hyperthermophilic Enzymes: Sources, Uses, and Molecular Mechanisms for Thermostability. Microbiol Mol Biol R 65:1–43.

Voss JE, Scally SW, Taylor NL, Atkinson SC, Griffin MDW, Hutton CA, Parker MW, Alderton MR, Gerrard JA, Dobson RCJ, et al. 2010. Substrate-mediated stabilization of a tetrameric drug target reveals Achilles heel in anthrax. J Biol Chem 285:5188–5195.

Wernegreen JJ. 2002. Genome evolution in bacterial endosymbionts of insects. Nat Rev Genet 3:850–861.

Wernegreen JJ. 2015. Endosymbiont evolution: predictions from theory and surprises from genomes. Ann Ny Acad Sci 1360:16–35.

Workneh F, Paetzold L, Rashed A, Rush CM. 2016. Population dynamics of released potato psyllids and their Bacteriliferous status in relation to zebra chip incidence in caged field plots. Plant Dis 100:1762–1767.

Wu D, Daugherty SC, Aken SEV, Pai GH, Watkins KL, Khouri H, Tallon LJ, Zaborsky JM, Dunbar HE, Tran PL, et al. 2006. Metabolic complementarity and genomics of the dual bacterial symbiosis of sharpshooters. PLoS Biol 4:188–14.

Yan Q, Sreedharan A, Wei S, Wang J, Pelz-Stelinski K, Folimonova S, Wang N. 2013. Global gene expression changes in Candidatus Liberibacter asiaticus during the transmission in distinct hosts between plant and insect. Mol Plant Pathol 14:391–404.

Ye J, Coulouris G, Zaretskaya I, Cutcutache I, Rozen S, Madden TL. 2012. Primer-BLAST: A tool to design target-specific primers for polymerase chain reaction. BMC Bioinform 13:134.

Yuan JS, Reed A, Chen F, Stewart CN. 2006. Statistical analysis of real-time PCR data. BMC Bioinform 7:85.

