## Supplementary data for "Genetic drift and genome reduction in the plant pathogen *Candidatus* Liberibacter solanacearum shapes a new enzyme in lysine biosynthesis"

### Supplementary Tables

**Supplementary Table i |** Comparison of kinetic constants between *C*LsoDHDPS and other bacterial homologues.

| Organism | $k_{cat}$ ($s^{-1}$) | $K_M^{pyruvate}$ (mM) | $K_M^{(S)\text{-ASA}}$ (mM) | $k_{cat}/K_M^{pyruvate}$ ($M^{-1}\ s^{-1}$) | $k_{cat}/K_M^{(S)\text{-ASA}}$ ($M^{-1}\ s^{-1}$) | $k_s$ |
|---|---|---|---|---|---|---|
| *Ca.* L. solanacearum (this work) | 6 | 0.012 | 0.033 | $5.0 \times 10^5$ | $1.8 \times 10^5$ | 0.002 |
| *E. coli* (Karsten 1997) | 188 | 0.19 | 0.12 | $9.9 \times 10^5$ | $1.6 \times 10^6$ | - |
| *M. tuberculosis* (Kefala et al. 2008) | 138 | 0.17 | 0.43 | $8.1 \times 10^5$ | $3.2 \times 10^5$ | - |
| *A. thaliana* (Griffin et al. 2012) | 93 | 1.0 | 0.09 | $9.3 \times 10^4$ | $1.0 \times 10^6$ | - |
| *T. maritima* (Pearce et al. 2006) | 5 * | 0.053 | 0.16 | $9.5 \times 10^4$ | $3.2 \times 10^4$ | - |
| *S. pneumoniae* (Dogovski et al. 2013) | 22 | 2.6 | 0.044 | $8.6 \times 10^3$ | $5.0 \times 10^5$ | - |
| *B. anthracis* (Domigan et al. 2009) | 76 | 0.43 | 0.18 | $1.8 \times 10^5$ | $4.2 \times 10^5$ | - |
| *N. meningitides* (Devenish et al. 2009) | 47 | 0.50 | 0.052 | $9.4 \times 10^4$ | $9.0 \times 10^5$ | - |
| *V. cholerae* (Gupta et al. 2018) | 34 | 0.14 | 0.08 | $2.4 \times 10^5$ | $4.3 \times 10^5$ | - |
| *C. jejuni* (Skovpen and Palmer 2013) | 76 | 0.35 | 0.16 | $2.2 \times 10^5$ | $4.8 \times 10^5$ | - |
| *K. pneumoniae* (Impey et al. 2020) | 290 | 1.2 | 0.34 | $2.4 \times 10^5$ | $8.5 \times 10^5$ | - |

* Data collected at 20 °C, far below this enzyme’s native temperature (~80 °C)

**Supplementary Table ii |** Data analysis statistics and collection parameters for the small angle X-ray scattering (SAXS) experiments.

| **Data analysis** | **apo-DHDPS** | **DHDPS + pyruvate** | **DHDPS + pyruvate/SSA** | **DHDPS + lysine** |
|---|---|---|---|---|
| *I*(0) ($cm^{-1}$) (Guinier analysis) | 0.11 ± 0.0002 | 0.0840 ± 0.0001 | 0.11 ± 0.0001 | 0.043 ± 0.0001 |
| $R_g$ (Å) (Guinier analysis) | 31.08 ± 0.30 | 32.93 ± 0.21 | 32.8 ± 0.25 | 31.5 ± 0.27 |
| *I*(0) ($cm^{-1}$) (*P*(*r*) analysis) | 0.11 | 0.08 | 0.11 | 0.04 |
| $R_g$ (Å) (*P*(*r*) analysis) | 31.08 | 32.93 | 32.8 | 31.5 |
| $D_{max}$ (Å) | 89.4 | 89.8 | 90.5 | 92.8 |
| Porod volume ($Å^{-3}$) | 164 000 | 214 000 | 172 000 | 171 000 |
| Molar mass (Porod volume, kDa) | 96 470 | 125 882 | 129 911 | 100 588 |
| Molar mass (SAXSMoW2*, kDa) | 113 650 | 123 248 | 101 176 | 124 742 |
| Tetrameric mass from sequence (kDa) | 132 000 | 132 000 | 132 000 | 132 000 |
| **Data collection parameters** | | | | |
| Instrument | Australian Synchrotron SAXS/WAXS beamline | | | |
| detector | PILATUS 1M (Dectris) | | | |
| wavelength (Å) | 1.0332 | | | |
| Maximum flux at sample | $8 \times 10^{12}$ photons per second at 12 keV | | | |
| Camera length (mm) | 1600 | | | |
| Q range ($Å^{-1}$) | 0.006-0.5 | | | |
| Exposure time | Continuous 1 second frame measurements | | | |
| Sample configuration | SEC-SAXS with co-flow | | | |
| Sample temperature (°C) | 12 | | | |

**Supplementary Table iii |** Data collection and refinement statistics for *C*LsoDHDPS with bound ligands. Values for the highest resolution shells are given in parentheses. Here, succinic semi-aldehyde is abbreviated to SSA.

| **Data collection statistics** | **DHDPS + lysine** | **DHDPS + pyruvate** | **DHDPS + pyruvate/SSA** |
|---|---|---|---|
| wavelength (Å) | 0.95369 | 0.95369 | 0.95369 |
| space group | C2 | C2 | C2 |
| unit cell parameters (a, b, c, Å) | 101.9, 133.5, 155.8 | 101.8, 133.6, 156.0 | 101.1, 132.9, 154.9 |
| resolution range (Å) | 45.2–2.01 (2.08–2.01) | 46.3–2.40 (2.49–2.40) | 45.8–1.93 (1.96–1.93) |
| observed reflections | 535,866 (53,764) | 317,539 (30,868) | 596,814 (26,666) |
| unique reflections | 135,620 (13,469) | 79,992 (7,961) | 151,510 (6,831) |
| mean $I/\sigma(I)$ | 13.5 (1.2) | 8.0 (1.0) | 7.8 (0.9) |
| completeness (%) | 99.9 (99.5) | 99.8 (99.8) | 99.5 (91.1) |
| $R_{merge}$ | 0.113 (0.553) | 0.344 (0.776) | 0.082 (1.25) |
| $R_{meas}$ | 0.131 (0.644) | 0.394 (0.900) | 0.11 (1.63) |
| $R_{pim}$ | 0.064 (0.315) | 0.191 (0.448) | 0.07 (1.11) |
| $CC_{½}$ | 0.954 (0.802) | 0.920 (0.702) | 0.998 (0.468) |
| Wilson *B*-factor ($Å^2$) | 31.4 | 36.9 | 36.7 |
| **Refinement statistics** | | | |
| $R_{factor}$ | 0.172 (0.239) | 0.199 (0.282) | 0.185 (0.400) |
| $R_{free}$ | 0.209 (0.281) | 0.229 (0.301) | 0.225 (0.430) |
| number of atoms | | | |
| non-hydrogen | 14,217 | 13,864 | 14,476 |
| macromolecules | 13,464 | 13,416 | 13,489 |
| solvent | 753 | 488 | 978 |
| protein residues | 1,776 | 1,776 | 1,776 |
| r.m.s.d. bonds (Å), angles (°) | 0.008, 1.0 | 0.022, 1.6 | 0.007, 1.0 |
| Ramachandran plot | | | |
| favoured, outliers (%) | 98.4, 0.3 | 97.6, 0.1 | 97.6, 0.2 |
| rotamer outliers (%) | - | 1.2 | - |
| clash score | 2.53 | 4.08 | 3.62 |
| PDB id | 7LVL | 7LOY | 8GEK |

**Supplementary Table iv** | Genes and primer sequences used in gene expression analysis.

| Gene | Accession number | Protein | Primer sequence |
|---|---|---|---|
| *dapA* | KJZ81861.1 | dihydrodipicolinate synthase | 5’-CACGGAGGTGTGGGTTGTAT-3’<br>3’-GTGCTTGACGATAATCCCCCT-5’ |
| *recA* | KJZ80672 | recombinase A | 5’-TACGCCCTTTGGGAAAACCA-3’<br>3’-TGCAGCTTGGCTCTCAAAGT-5’ |
| *Rpb* | KJZ81365.1 | DNA-directed RNA polymerase beta subunit | 5’-CCACAAGATACAATCGCCGC-3’<br>3’-GCGGGAAAAGGTTTACTGCG-5’ |

**Supplementary Table v** | Primers used to determine *Ca*. L. solanacearum titres in DNA samples and to serve as internal psyllid controls.

| Primer | Primer sequence |
|---|---|
| *C*LsoF | 5’-GTCGAGCGCTTATTTTTAATAGGA-3’ |
| *C*Lso16SF | 5’-ATACCGTATACGCCCTGAGAAG-3’ |
| *C*Lso16SRI | 3’-CGTAGCCTTGGTAGGCATT-5’ |
| ITS2F | 5’-AAGCGACGTGTGGAAGAACC-3’ |
| ITS2R | 3’-GTTGTGTGTGTCCGGGGAAG-5’ |
| COX-F | 5’-CGTCGCATTCCAGATTATCCA-5’ |
| COX-R | 3’-CAACTACGGATATATAAGAGCCAA AAC-5’ |

### Supplementary Figures

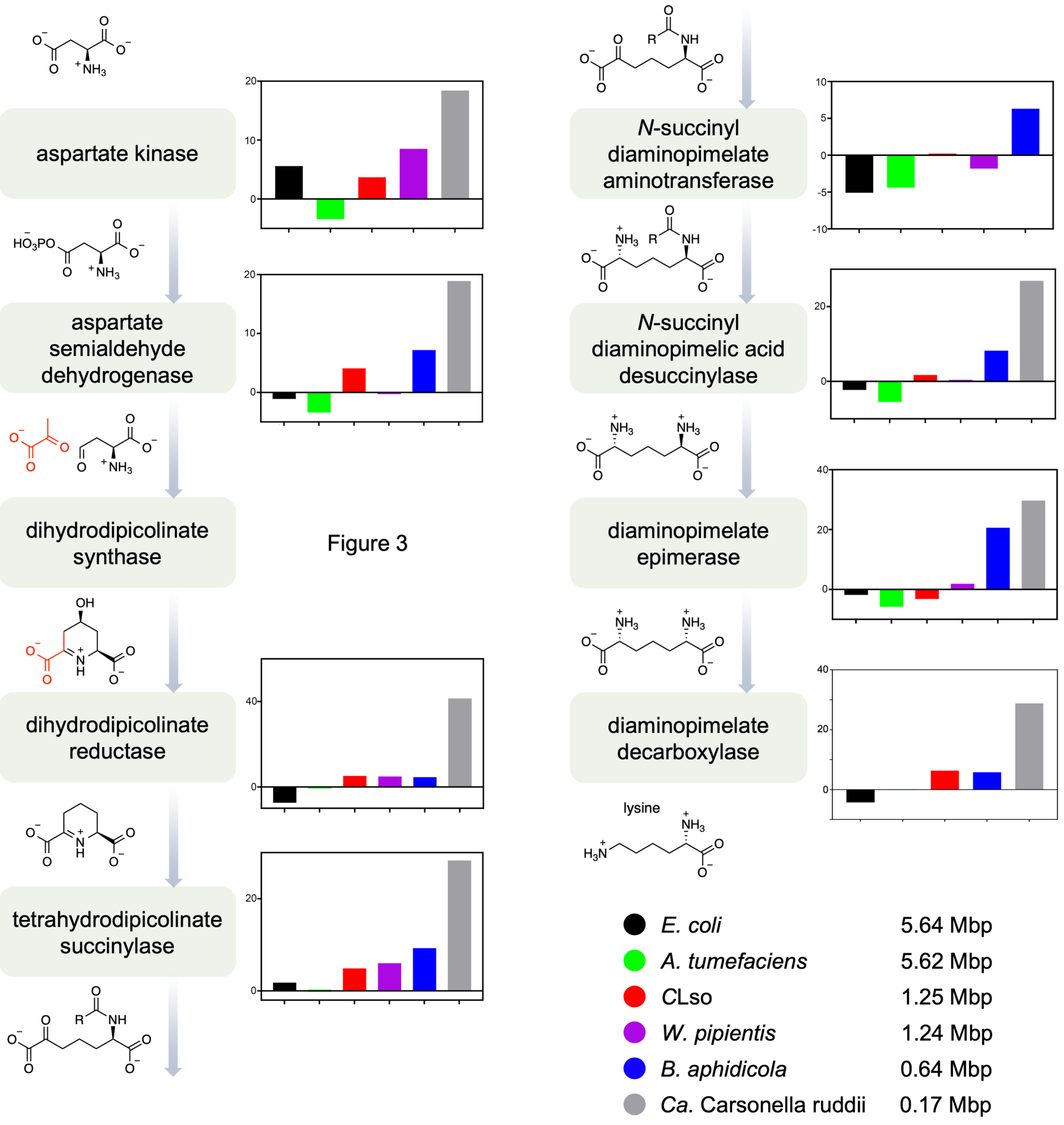


**Supplementary Figure i |** Aggregation propensity of enzymes in the lysine biosynthesis pathway highlight the trend of increased aggregation as the genome size is reduced. An increased positive $Na^4vSS$ score means an increase in the aggregation propensity for the protein, whereas a negative number suggests a decreased aggregation propensity.

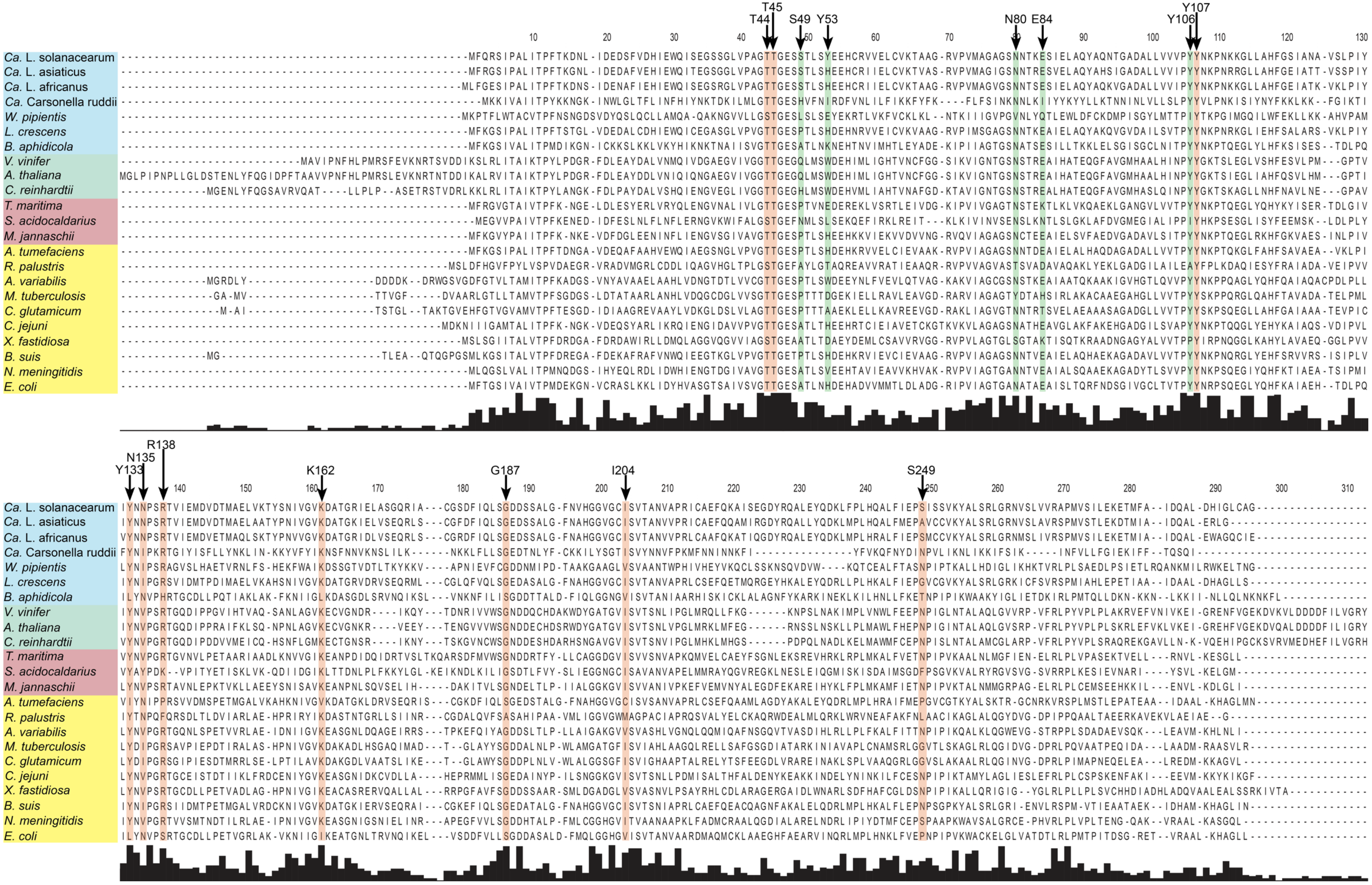


**Supplementary Figure ii | Sequence alignments of *C*LsoDHDPS (top sequence) with representative bacterial and plant homologues.** Blue resents sequences from reduced genome bacteria, green from plant/algal, pink from extremophile bacteria, and yellow from mesophilic bacteria. Residues shaded in green represent those that fin lysine in the allosteric site, and those shaded on orange are active site residues that interaction with the substrates

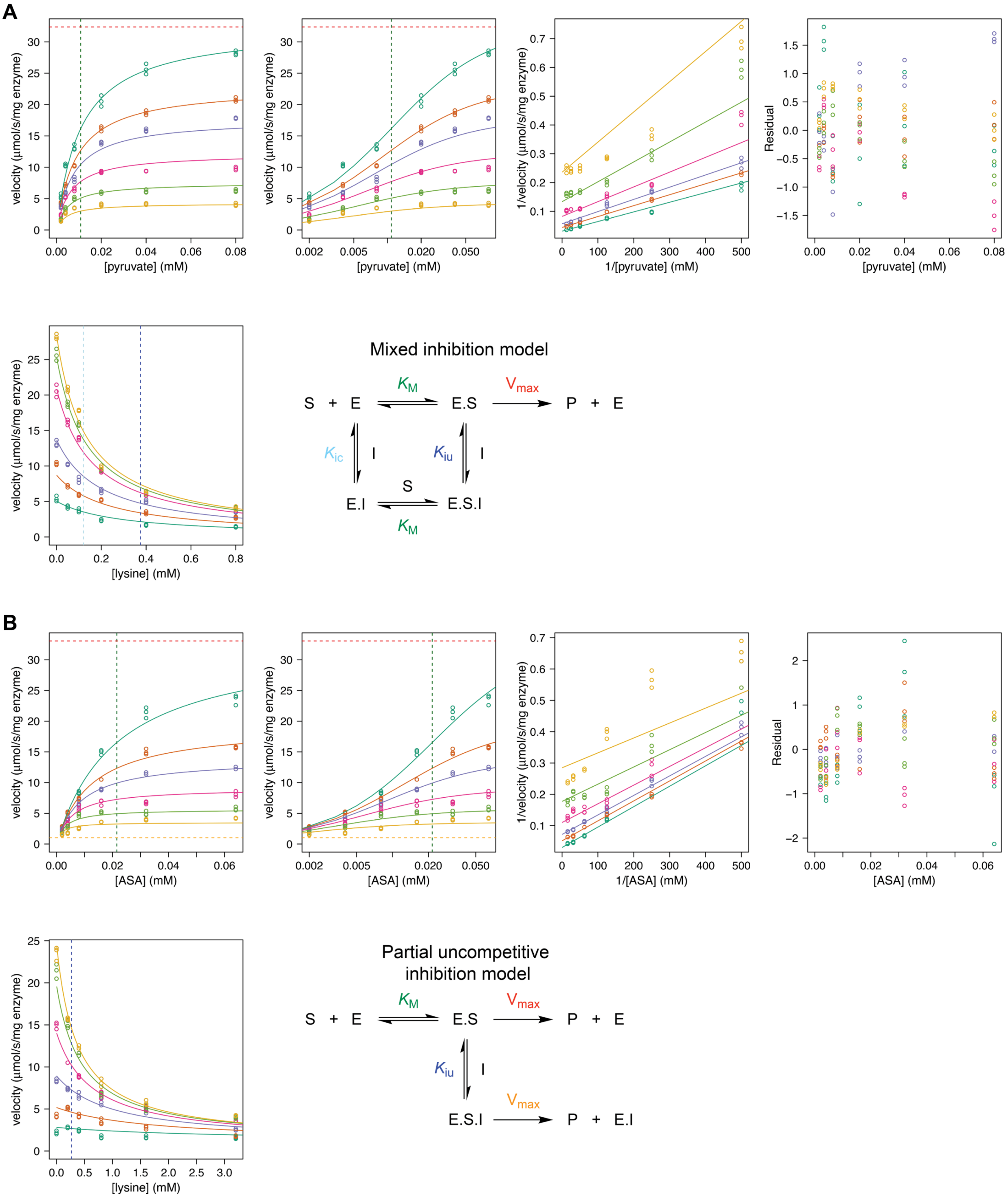


**Supplementary Figure iii | *CLso*DHDPS is inhibited by lysine. A** is with respect to pyruvate at different lysine concentrations fitted to a mixed model inhibition. From left to right (first row) the plots include a direct plot, the direct plot with the substrate concentration on the log scale, the Lineweaver Burk plot, and the residuals of the fit. The second row displays a direct plot with respect to lysine at different concentrations of pyruvate as well as schematic representations of the mixed inhibition model. **B** is with respect to *S*-aspartate-β-semialdehyde (ASA) at different lysine concentrations fitted to a partial uncompetitive inhibition model. From left to right (first row) the plots include a direct plot, the direct plot with the substrate concentration on the log scale, the Lineweaver Burk plot, and the residuals of the fit. The second row displays a direct plot with respect to lysine at different concentrations of *S*-aspartate-β-semialdehyde (ASA) as well as a schematic representation of the partial uncompetitive inhibition models. In both A and B, the red dotted line shows the fitted $V_{max}^{app}$, the green dotted line is the $K_M^{app}$, and the blue dotted line is the $K_i^{lysine}$. All fits and plots were generated using R.

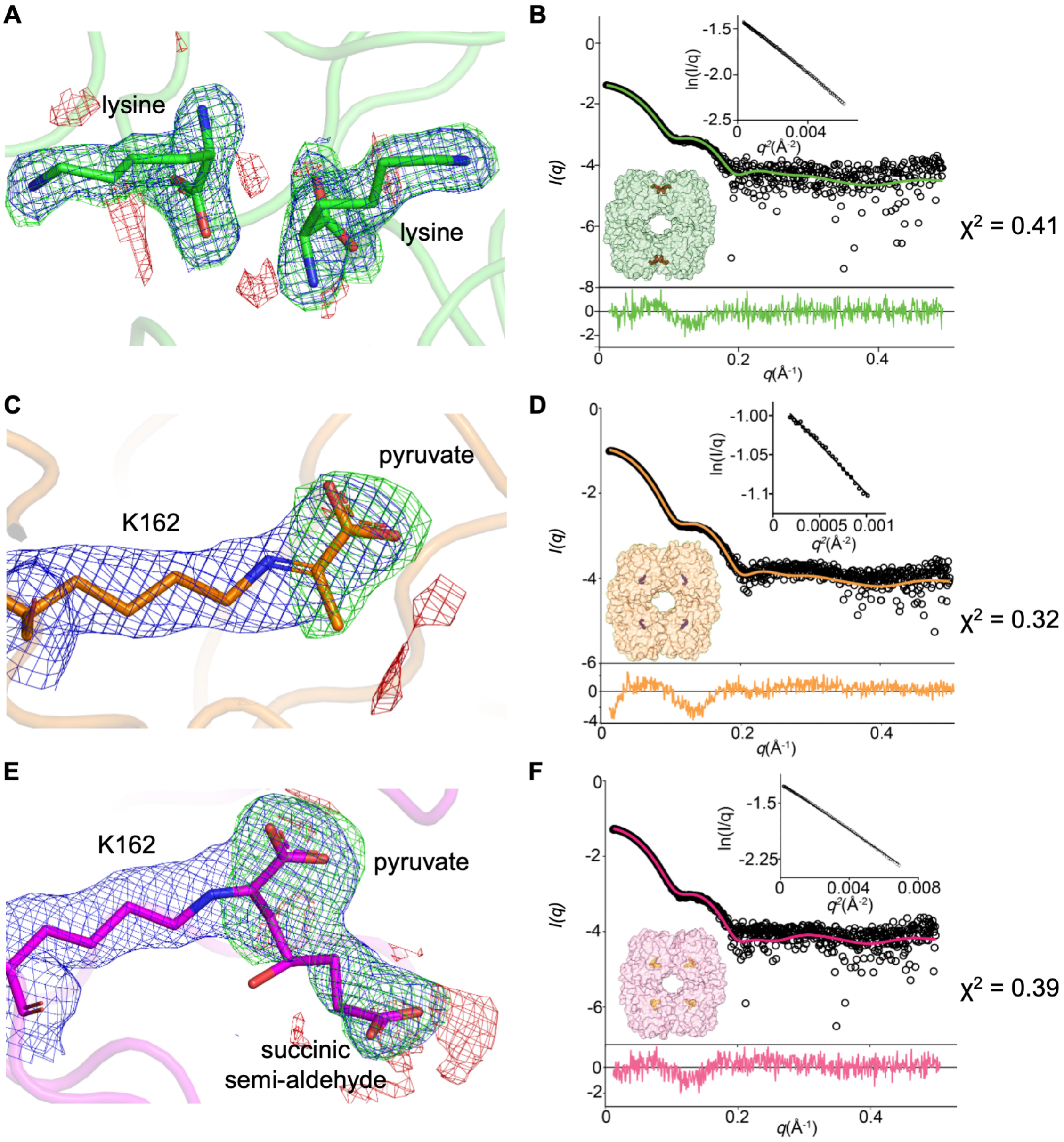


**Supplementary Figure iv | Omit maps and small angle X-ray scattering analysis of *C*LsoDHDPS with ligands. A, C** and **E.** Omit maps around the ligands bound to *C*LsoDHDPS [**A**, lysine (green); **C**, pyruvate (orange); and **E**, pyruvate + succinic semi-aldehyde (magenta)]. For all electron density maps, the 2Fo-Fc map is set to 1 σ (blue mesh), and the difference density Fo-Fc map is set to 3 σ (green mesh) and -3 σ (red mesh). **B, D** and **F.** Experimental scatter (o) of *C*LsoDHDPS with ligands (**B**, lysine, 10 mM; **D**, pyruvate 5 mM; and **E**, pyruvate + succinic semi-aldehyde, 5 mM) fitted to the back calculated scatter from the equivalent ligand bound tetrameric crystal structures of *C*LsoDHDPS. The residuals for the fit are shown (bottom) along with the surface structure of *C*LsoDHDPS showing ligands (red sticks, bottom left) and the Guinier plot (inset top right). The $\chi^2$ value for each fit is indicated on the right.

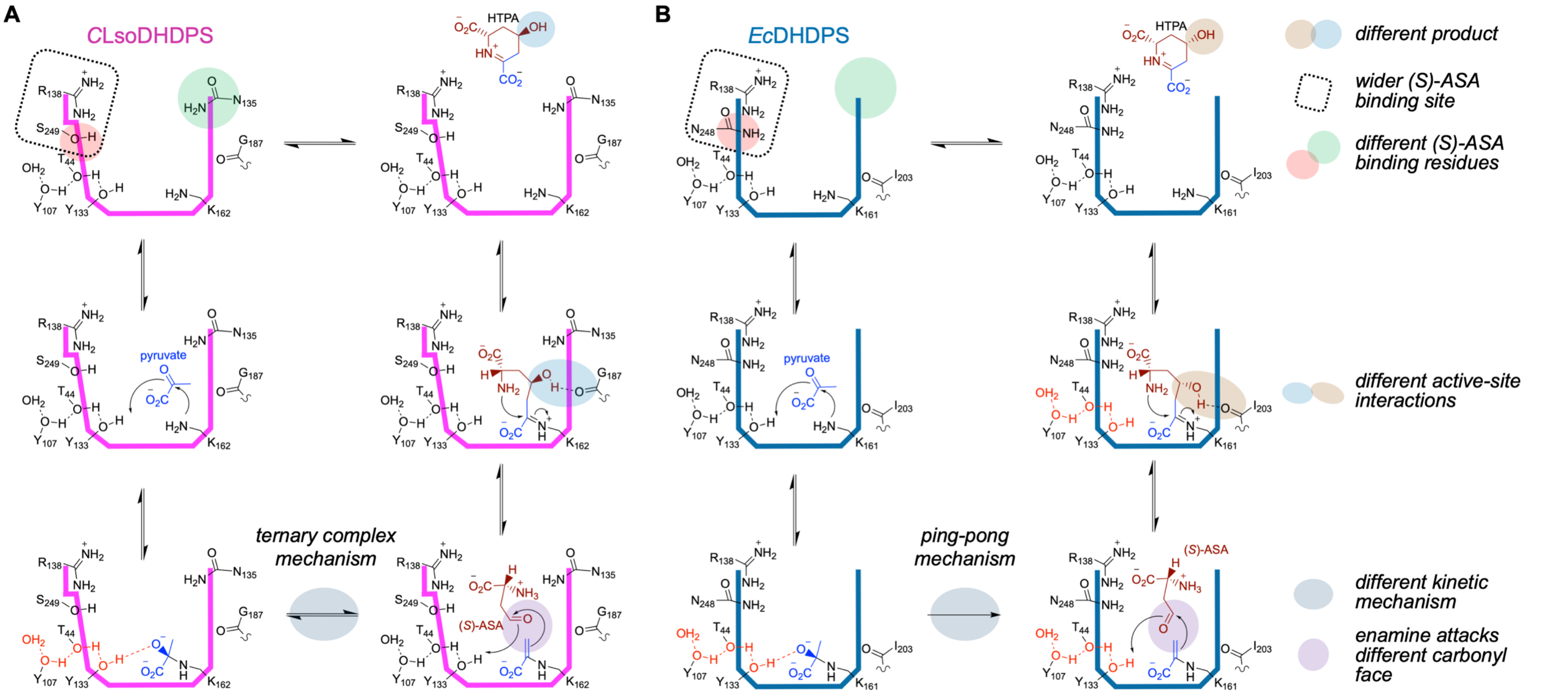


**Supplementary Figure v | Omit maps and small angle X-ray scattering analysis of *C*LsoDHDPS with ligands. A** and **B.** Schematic highlighting the differences in the proposed mechanisms for *C*LsoDHDPS and *Ec*DHDPS, as per **Figures 7** and **8**.
